# Jag1 represses Notch activation in lateral supporting cells and inhibits an outer hair cell fate in the medial compartment of the developing cochlea

**DOI:** 10.1101/2024.02.02.577075

**Authors:** Sandra de Haan, Agustin A. Corbat, Christopher R. Cederroth, Lisa G. Autrum, Simona Hankeova, Elizabeth C. Driver, Barbara Canlon, Matthew W. Kelley, Emma R. Andersson

## Abstract

Notch signaling regulates both inner and middle ear morphogenesis and establishes a strict pattern of sensory cells in the organ of Corti in the mammalian cochlea. Patients with Alagille syndrome have impaired Notch signaling (∼94% with *JAG1* mutations) resulting in sensorineural and conductive hearing loss. Here, we investigate the function of Jag1-mediated Notch activation in cochlear patterning and signaling using the Jag1 “Nodder” (*Jag1*^*Ndr/Ndr*^) mouse model of Alagille syndrome. *Jag1*^*Ndr/Ndr*^ mice exhibited severe vestibular and auditory deficits and a dose-dependent increase in ectopic inner hair cells and a reduction in outer hair cells. Single cell RNA sequencing of the organ of Corti demonstrated a global dysregulation of genes associated with inner ear development and deafness. Analysis of individual cell types indicated a novel role for Jag1 in repressing Notch activation in lateral supporting cells and revealed a function for Jag1 in gene regulation and development of outer hair cells. Additionally, “outer hair cell-like” SLC26A5 (Prestin) positive cells were present in the medial compartment and pillar cell region of *Jag1*^*Ndr/Ndr*^ mice and exhibited location-dependent expression of the inner hair cell fate-regulator *Tbx2*, revisiting the potency of *Tbx2* in driving inner hair cell commitment in “outer hair cell-like” cells in the Jag1-mutant IHC compartment. Our data reveals a novel role for Jag1 in repressing Notch activation in lateral supporting cells and highlights involvement for Notch signaling in inner versus outer hair cell specification and positioning.

## Introduction

Proper auditory function requires a precisely ordered mosaic pattern of mechanosensitive inner hair cells (IHCs) and outer hair cells (OHCs) as well as several subtypes of associated supporting cells (SCs) within the organ of Corti. Organ of Corti formation is controlled by a series of cell fate decisions mediated by Notch signaling, a conserved signaling pathway acting through membrane bound ligand – receptor interactions. Two distinct Notch-mediated signaling mechanisms drive pattern formation: lateral induction, by which the Notch ligand JAG1 activates Notch and forms the prosensory domain containing progenitor cells that will give rise to the variety of cell types in the organ of Corti, and lateral inhibition, by which the Notch ligands DLL1 and JAG2 repress the hair cell (HC) fate in contacting cells. Both signaling modes rely on trans-signaling events, in which Notch receptors are activated by Notch ligands on juxtaposed cells.

In humans, *JAG1* mutations are associated with the congenital disorder Alagille syndrome (ALGS)^1–4^. Although ALGS is best known as a liver disorder, patients also exhibit hearing deficits including conductive and sensorineural hearing loss, which are due to middle ear bone malformations or sensory/neural cell deficits, respectively^5^. Balance can also be affected in patients, as temporal bone defects including loss of the posterior semicircular canal have been reported^6^. Finally, patients with *JAG1* mutations and hearing deficits but no liver involvement have also been reported^7^. Understanding how *Jag1* insufficiency impacts ear development in a model of ALGS could thus provide deeper insights into hearing loss in patients.

The best-described Notch loss-of-function phenotype, induced by *Notch1* deletion or by a gamma-secretase inhibitors, is an excess of IHCs and OHCs at the expense of SCs, which arises when the HC fate is not repressed by Notch activation in SCs^8^ (for an overview of Notch defective models, see^8^ and **Supp. Table 1**). In contrast with Jag2 and Dll1, which are expressed only in developing HCs, Jag1 is expressed during the entire time window of cochlear development: from early formation of the prosensory domain, then alongside *Jag2* and *Dll1* during cell fate acquisition, and finally during cochlear maturation, when its expression is maintained in SCs. Defects in Jag1 result in supernumerary IHCs but fewer OHCs, and subsequent hearing deficits, as exemplified by mice with mutations in *Jag1* ^9–12^ (**Supp. Table 1, 2**). More recently, conditional deletion of Jag1 in SCs was shown to result in the absence of Hensen’s cells (HeCs)^10,13^. Previous reported models, however, are limited to conditional deletion and heterozygous loss of function of *Jag1* due to late embryonic lethality of *Jag1* germline mutants. It therefore remains elusive how *Jag1* insufficiency affects the development of individual SCs and HCs, in particular IHCs and OHCs, both at transcriptomic level and in cellular patterning.

Here, we investigate the role of Jag1-mediated Notch activation in cochlear patterning and signalling, using the “Nodder” (*Jag1*^*Ndr/Ndr*^) mouse model for ALGS, which is, in contrast to previously reported models, viable in the homozygous condition. *Jag1*^*+/Ndr*^ and *Jag1*^*Ndr/Ndr*^ mice exhibited severe patterning defects, including a dose-dependent increase in ectopic IHCs, and a reduction in OHCs, with a concomitant increase and decrease in associated medial and lateral SCs, respectively. Transcriptomic analyses of *Jag1*^*Ndr/Ndr*^ cochleae demonstrated a global dysregulation of genes associated with inner ear development and deafness. Single cell RNA sequencing (scRNAseq) furthermore revealed an upregulation of Notch target genes in *Jag1*^*Ndr/Ndr*^ lateral SC populations, indicating that Jag1 limits Notch signalling in lateral SCs. In addition, there was major transcriptomic dysregulation in OHCs, revealing a function for *Jag1* in HC subtype gene regulation and development. Finally, we identified ectopic SLC26A5 (Prestin) positive “outer hair cell-like” (OHC-like) cells in the IHC and pillar cell (PC) compartments, with OHC-like cells in the IHC compartment expressing the IHC fate regulator *Tbx2*. These findings indicate that *Tbx2* is not sufficient to drive IHC commitment in OHC-like cells in the *Jag1*-mutant IHC compartment and highlights a novel role for Notch signalling in IHC vs OHC fate acquisition.

## Results

### Adult *Jag1*^*Ndr/Ndr*^ mice exhibit major vestibular and auditory defects

To assess the impact of the *Jag1* Nodder mutation on functional characteristics of the inner ear, we first assessed balance and hearing function in *Jag1*^*Ndr/Ndr*^ mice (“Nodder” mice), which recapitulate major hallmarks of ALGS^14^. As reported previously, but not shown, *Jag1*^*Ndr/Ndr*^ mice exhibit a repetitive up-down head movement, with emphasis on head tossing upwards towards the shoulders (**Fig 1a, 1s panel, Supp. video 1**). Analysis of the vestibular system at postnatal day (P)45-P60 demonstrated an absence of the posterior semicircular canal, which detects head-tilting towards the shoulders **(Fig 1b**). In addition to the distinctive head nodding behaviour, *Jag1*^*Ndr/Ndr*^ exhibit circling behaviour and are hyperactive (**Fig 1c-d**).

**Figure 1:**
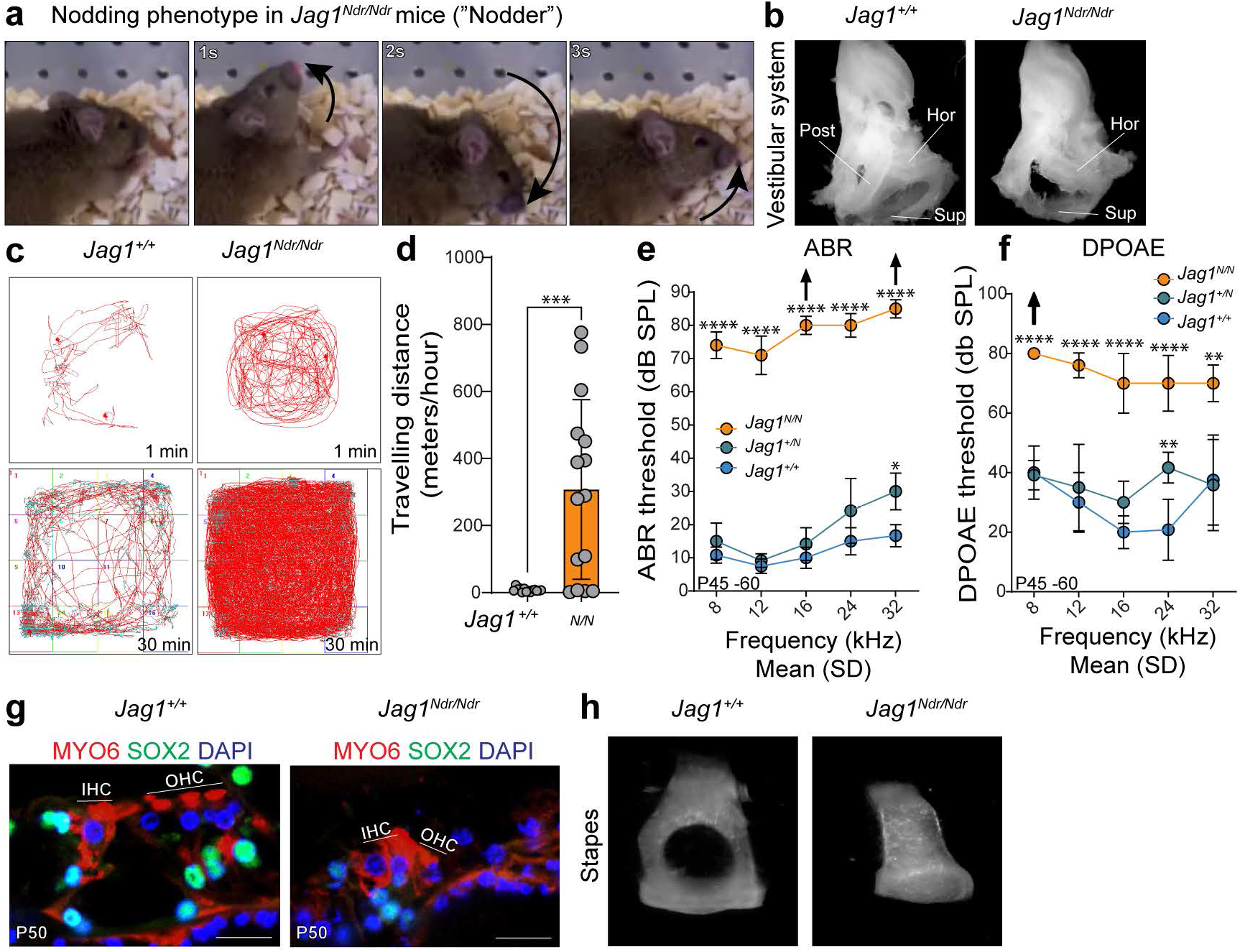
Adult *Jag1*^*Ndr/Ndr*^ mice exhibit major vestibular and auditory defects. **a)** Timelapse images of head nodding behaviour of *Jag1*^*Ndr/Ndr*^ mice, showing movement of the head upwards and towards the shoulders. **b)** *Jag1*^*+/Ndr*^ and *Jag1*^*Ndr/Ndr*^ vestibular systems, showing the absence of the posterior semicircular canal in *Jag1*^*Ndr/Ndr*^ mice. **c,d)** Open Field Test tracking for 1 or 30 mins, demonstrating that **(c)** *Jag1*^*Ndr/Ndr*^ mice run in circles and **(d)** are hyperactive, as measured by meters travelled per hour (distance travelled = 307.7±267.8 meters/hour (m/h) in *Jag1*^*Ndr/Ndr*^ versus 7.5±5.8 m/h in *Jag1*^+/+^, p-value <0.001, mean ± SD). **e,f)** Auditory measurements indicate hearing loss with increased ABR thresholds **(e)** and DPOAE thresholds in *Jag1*^*Ndr/Ndr*^ mice **(f)**. Arrows indicate no auditory response was measured at the highest sound stimulus. **g)** HC and SC phenotype at adult stage (P50) indicates a disorganised pattern of *Jag1*^*Ndr/Ndr*^ HCs (MYO6^+^ cells, red) and SCs (Sox2^+^ cells, green). **h)** *Jag1*^*Ndr/Nd*^ display malformed, columnar stapes. Data are mean with standard deviation (SD); each datapoint in **(d)** represents measurements from one mouse; scale bar represents 20 µm; *p-value <0.05; **p-value <0.01; ***p-value <0.001, ****p-value <0.0001; unpaired t-test for travelling distance (**d**) and one-way ANOVA with Bonferroni correction for auditory measurements, n=6 per genotype (**e, f**); Post=posterior; Hor=horizontal; Sup= Superior; ABR=Auditory Brain Stem Response; DPOAE=Distortion Product Otoacoustic Emission; min=minutes; HC=hair cell; SC=supporting cell; IHC=inner hair cell; OHC=outer hair cell.

Hearing loss is reported in ALGS^5,7,15,16^, and in heterozygous mutant and *Jag1* conditional loss-of-function mice^9–12^ (**Supp. Table 2**). We therefore assessed hearing function by measuring auditory brain stem response (ABR) and distortion product otoacoustic emissions (DPOAE) in adult (P45-P60) *Jag1*^*+/+*^, *Jag1*^*+/Ndr*^ and *Jag1*^*Ndr/Ndr*^ mice. *Jag1*^*Ndr/Ndr*^ mice had severe hearing deficits with ABR thresholds above 70 decibels (dB SPL) along all measured frequencies, while *Jag1*^*+/Ndr*^ mice had a mild hearing deficit only at higher frequencies (**Fig 1e**). Additionally, DPOAE thresholds were elevated among all frequencies in *Jag1*^*Ndr/Ndr*^ mice, indicating reduced cochlear amplification and OHC dysfunction (**Fig 1f**). Histological examination of the adult cochlea demonstrated the presence of some Myosin6^+^ (MYO6) HCs, although in a less organised pattern **(Fig 1g**), providing a possible basis for sensorineural hearing loss. Finally, as both sensorineural and conductive hearing loss are reported in Alagille syndrome, we assessed the middle ear bones to determine whether conductive function is compromised and found that *Jag1*^*Ndr/Ndr*^ mice exhibited malformed stapes (**Fig1h**). In sum, *Jag1*^*Ndr/Ndr*^ mice exhibit profound balance and hearing defects, with vestibular and middle ear bone malformations, and HC defects, which together recapitulate the mixed hearing loss in patients with Alagille syndrome.

### Inner ear development and hearing genes are dysregulated in the *Jag1*^*Ndr/Ndr*^ organ of Corti

*Jag1* loss of function results in supernumerary IHCs, a reduction in OHCs^11,17,18^, and SC survival defects^9,10,19^, and disorganised HCs were present in both adult *Jag1*^*Ndr/Ndr*^ mice (**Fig1g**) and at postnatal day P5 (**Fig2a**). Therefore, to investigate the consequences of JAG1 insufficiency in individual cell types in *Jag1*^*Ndr/Ndr*^ mice, we characterised transcriptional changes in the *Jag1*^*Ndr/Ndr*^ organ of Corti by scRNAseq. We collected cells from the organ of Corti of *Jag1*^*+/+*^ and *Jag1*^*Ndr/Ndr*^ mice at P5, when individual cell types of the organ of Corti have differentiated and are undergoing maturation^20^. After quality control and filtering, 11,416 cells were analyzed (**Fig 2a-b**). Dimensionality reduction using UMAP and clustering identified 21 cell types/states, including different cell types within, medial, and lateral to the organ of Corti, as well as periotic mesenchyme and glial populations (**Fig2b-c, Supp.Table3**).

**Figure 2:**
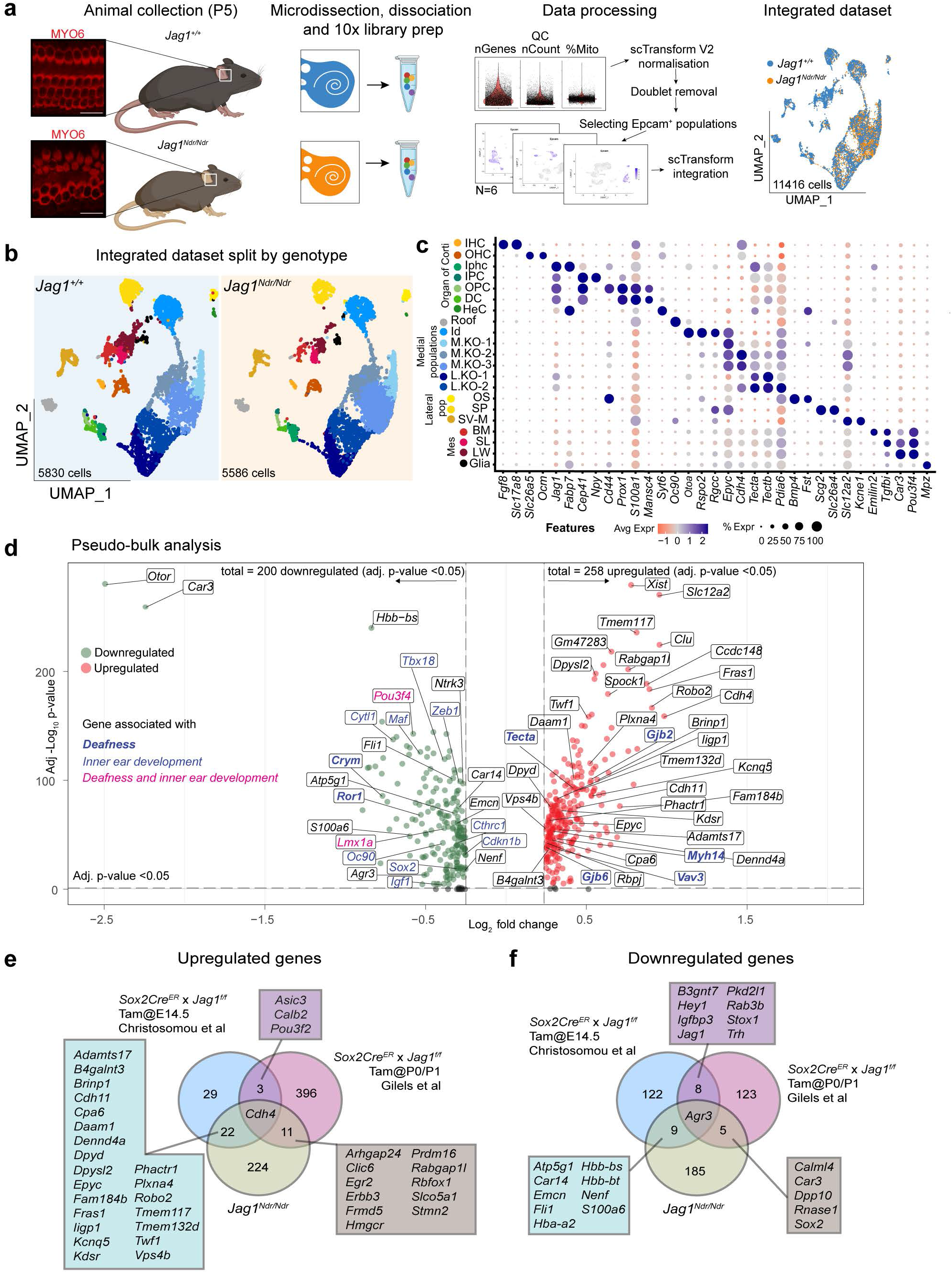
Inner ear development and hearing genes are dysregulated in the *Jag1*^*Ndr/Ndr*^ organ of Corti. **a)** Summary image of scRNAseq workflow. **b)** UMAP of *Jag1*^*Ndr/Ndr*^ and *Jag*^*+/+*^ integrated datasets (n=3 per genotype), split by genotype. **c)** Dot plot showing marker genes used to identify and annotate different cell populations. **d)** Volcano plot showing pseudo-bulk downregulated genes (left, green) and upregulated genes (right, red) between *Jag1*^*Ndr/Ndr*^ and *Jag*^*+/+*^, highlighting genes associated with deafness (blue, bold), inner ear development (blue), or both (magenta). **e,f)** Venn diagram showing the overlap between pseudo-bulk analysis identified **e)** up-regulated and **f)** down-regulated genes in the *Jag1*^*Ndr/Ndr*^ dataset compared with genes previously reported to be up- or down-regulated in Jag1 defective models. HeC=Hensen’s cell; Id=interdental; OS=outer sulcus; M.KO=medial Köllikers organ; L.KO=lateral Kölliker’s organ; SP=spiral prominence; SV-M=stria vascularis marginal; BM=basement membrane; SL=spiral limbus; LW=lateral wall.

To compare the scRNAseq with previously published *Jag1*-related bulk RNAseq datasets^9,10^, we first performed a pseudo-bulk analysis of the *Jag1*^*Ndr/Ndr*^ and *Jag1*^*+/+*^ populations. Differential gene expression analysis for the cochlear epithelium dataset revealed 458 dysregulated genes (avg. Log 2 FC >0.25, or <-0.25, and adj. p-value <0.05), including genes linked to deafness (https://www.ebi.ac.uk/gwas/efotraits/EFO_0004238, **Fig 2d, Supp. Table 4**, such as *Pou3f4*^21,22^, Crym^23^, *Ror1*^24^ *and Lmx1a*^25^), and contributing to the GO terms “Inner ear development” and “Inner ear morphogenesis” (**Fig 2d, Supp. Table 4**, GO:0048839 and GO:0042471, such as *Sox2*). *Jag1*^*Ndr/Ndr*^ dysregulated genes overlapped with genes dysregulated upon *Jag1* silencing in SCs^9,10^, specifically upregulation of the medial SC marker *Cdh4*^26^ in all models, and downregulation of *Agr3*, which is most strongly expressed in lateral SCs^27^ (**Fig 2e-f**). Therefore, the *Jag1*^*Ndr/Ndr*^ inner ear exhibits dysregulated gene signature associated with inner ear development, morphogenesis, and hearing, and recapitulates aspects of published *Jag1* loss-of-function transcriptomic data.

### *Jag1*^*Ndr/Ndr*^ mice exhibit a medial boundary defect, a reduction in lateral SCs and HeC ablation

As the pseudo bulk analysis demonstrated an overlap in dysregulated genes between *Jag1*^*Ndr/Ndr*^ mice and models with conditional deletion of *Jag1* in SOX2-expressing SCs (**Fig 2e-f**), we next assessed SC patterning and abundance, as well as transcriptional changes for each of the SC subtypes in *Jag1*^*Ndr/Ndr*^ mice.

*Jag1*^*Ndr/Ndr*^ mice exhibited a duplicated row of IHCs, accompanied by a duplication of FABP7^+^ IPhCs (**Fig 3a-b)**, indicating a medial boundary defect that includes a proportional duplication of both IHCs and IPhCs. Differential gene expression analysis identified 40 up- and 42-downregulated genes in *Jag1*^*Ndr/Ndr*^ versus *Jag1*^*+/+*^ IPhCs, with pathway dysregulation similar to the pseudobulk analyses (**Fig3c, Supp. Table 5,6**). To study cell identity, we manually curated a list of 200 genes specific for each cell type based on gene expression in the *Jag1*^*+/+*^ dataset, resulting in signature genes for each cell type (**Supp. Table 7**). Downregulated genes in *Jag1*^*Ndr/Ndr*^ IPhCs did not include IPhC signature genes, indicating that *Jag1*^*Ndr/Ndr*^ IPhC identity was not affected by JAG1 insufficiency.

**Figure 3:**
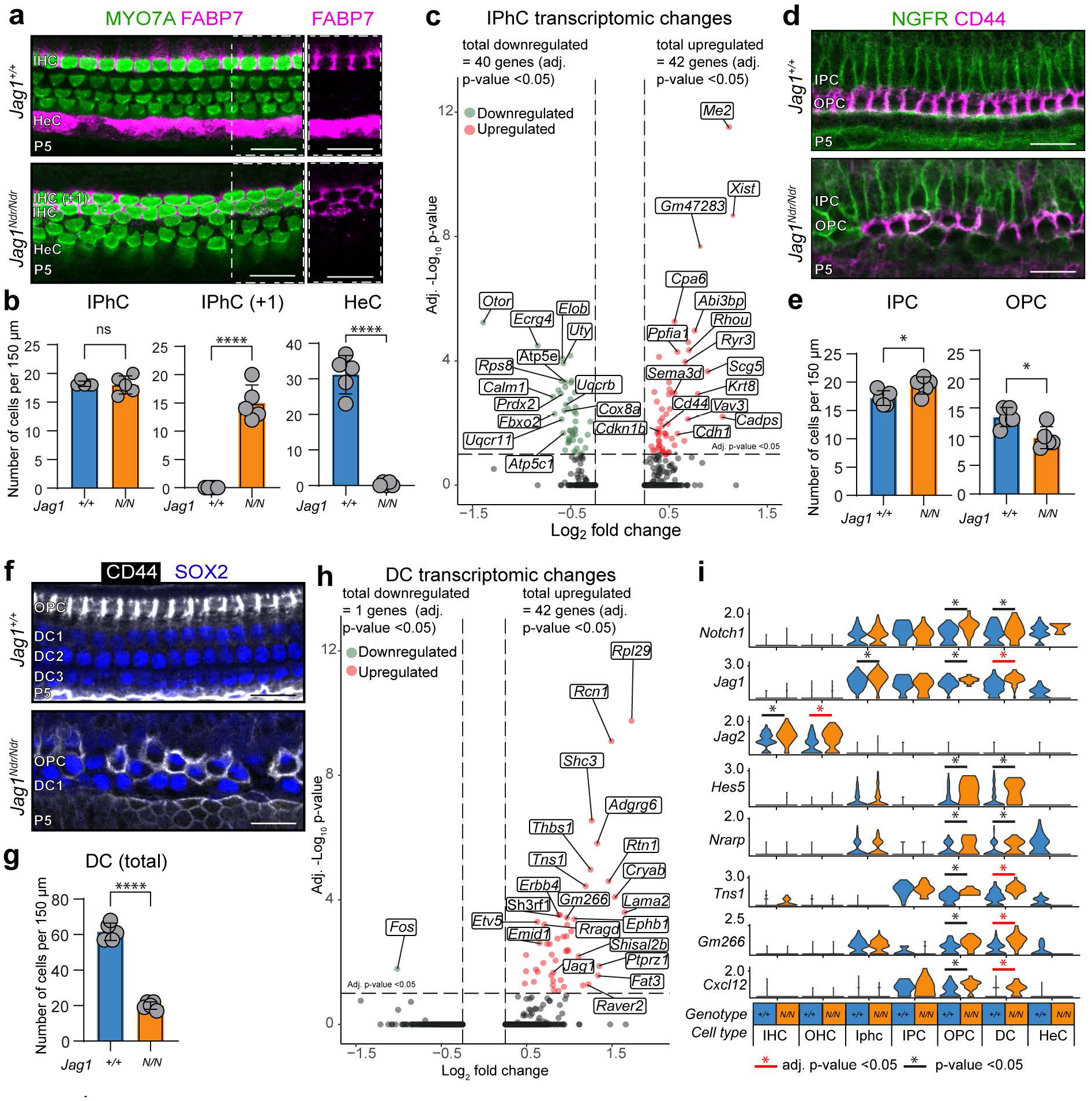
P5 *Jag1*^*Ndr/Ndr*^ mice exhibit supernumerary IPhCs, fewer lateral SCs, HeC ablation, and increased Notch activation in lateral SCs. **a)** HC, IPhC and HeC phenotype at P5, showing a duplication of IHCs (MYO7A, green) and IPhCs (FABP7, magenta) in *Jag1*^*Ndr/Ndr*^ cochleae. Right panel shows FABP7 alone, highlighting the lack of FABP7 signal in the lower *Jag1*^*Ndr/Ndr*^ HeC region. **b)** Quantification of IPhCs, supernumerary IPhCs (IPhC +1), showing18.0±1.6 IPhCs cells per 150 µm in *Jag1*^*Ndr/Ndr*^ compared to 18.2±0.5 in *Jag1*^*+/+*^, p-value=ns and 15.0±3.2 with supernumerary IPhCs cells (“IPhC +1”) per 150 µm in *Jag1*^*Ndr/Ndr*^ compared to 0.0±0.0 in *Jag1*^*+/+*^, p-value =<0.0001, mean ± SD, and 0.4±0.6 HeC cells per 150 µm in *Jag1*^*Ndr/Ndr*^ compared to 31.2±5.4 in *Jag1*^*+/+*^, p-value = <0.0001, mean ± SD, **c)** Volcano plot depicting up- and down-regulated genes (adj. p-value <0.05) in P5 *Jag1*^*Ndr/Ndr*^ IPhCs. **d**) PC phenotype, showing IPCs (NGFR, green) and OPCs (CD44, magenta). **e)** P5 quantification of IPCs and OPCs in the mid region, demonstrating an increase in IPCs, and a decrease in OPCs, with 19.4±1.5 IPC cells per 150 µm in *Jag1*^*Ndr/Ndr*^ compared to 17.2±1.3 in *Jag1*^*+/+*^, p-value =0.05, 9.8±1.9 OPC cells per 150 µm in *Jag1*^*Ndr/Ndr*^ compared to 13.4±1.6 in *Jag1*^*+/+*^, p-value =<0.05, mean ± SD. **f)** DC phenotype: staining for DCs (SOX2, blue) and OPCs (CD44, white) demonstrates a reduction of DCs lateral from OPCs. **g)** Quantification of DCs demonstrates a decrease in DCs in *Jag1*^*Ndr/Ndr*^ mice, with 20.0±2.1 DC cells per 150 µm in *Jag1*^*Ndr/Ndr*^ compared to 61.6±4.9 in *Jag1*^*+/+*^, p-value = <0.0001, mean ± SD **h)** Volcano plot depicting up- and down-regulated genes (adj. p-value <0.05) in *Jag1*^*Ndr/Ndr*^ DCs. **i)** Violin plots depicting the expression of Notch components per genotype and cell type, indicating increased Notch activation in OPCs and DCs. **j)** Volcano plots depicting expression of genes identified by Castel et al^35^ to contain Rbpj binding site, showing up-regulation of these genes in lateral SCs, but not HeCs, in *Jag1*^*Ndr/Ndr*^ mice. n=5 or more per genotype for phenotype characterisation and each dot represents one animal; red asterisks indicate significant upregulation (adj. p-value <0.05); black asterisks indicates potential upregulation (p-value <0.05); *p-value <0.05; **p-value <0.01; ***p-value <0.001, ****p-value <-0.0001, unpaired t-test).

FABP7 (a Notch target gene also known as *Blbp*^28^) is a marker of IPhCs, as well as HeCs^10,29^ which are present lateral to the OHCs in the *Jag1*^*+/+*^ organ of Corti (**Fig 3a, right panel**). However, FABP7 expression was almost completely absent in the lateral compartment along the entire length of the cochlea in *Jag1*^*Ndr/Ndr*^ mice **(Fig 3a-b**), and we only sporadically found a FABP7^+^ HeC (**Supp. Fig 1a-b**). FABP7 absence in the lateral compartment could reflect Notch silencing in HeCs, rather than a de facto loss of HeCs. However, 43 *Jag1*^*+/+*^ and only 3 *Jag1*^*Ndr/Ndr*^ HeCs were retrieved in the scRNA seq (**Fig 2b, Supp. Table 8**), corroborating that HeCs are depleted in *Jag1*^*Ndr/Ndr*^ mice.

Next, we assessed PCs, a specialised SC subtype located between IHCs and OHCs whose differentiation and maintenance are Notch-independent^30^. PCs express Nerve Growth Factor Receptor (NGFR, also known as p75^ntr^)^31–33^ and outer PCs (OPCs) begin to express CD44 between P0 and P7^34^. There was an increase in NGFR^+^ PCs and a decrease in CD44^+^ OPCs in the mid region at P5 (**Fig3d-e**), while IPC and OPC numbers were similar in the base region **(Supp. Fig 1c-d)**, indicating that PC development may be delayed in *Jag1*^*Ndr/Ndr*^ mice. We therefore assessed the PC phenotype at birth. As expected, *Jag1*^*+/+*^ mice exhibited one row of NGFR^+^ IPC accompanied by one row of CD44^+^ OPC. However, *Jag1*^*Ndr/Ndr*^ mice showed a delay in PC development, displaying a more widespread NGFR^+^ signal reaching two to three cell layers, and almost completely lack CD44^+^ cells, mimicking a less mature phenotype that is typically observed around E17.5 in *Jag1*^*+/*+^ mice (**Supp. Fig 1c**). Despite the developmental delay observed in IPCs and OPCs, only minor transcriptomic differences were detected in IPCs and OPCs between *Jag1*^*Ndr/Ndr*^ and *Jag1*^*+/+*^ animals, with only one gene, *Rpl29*, being upregulated in OPCs (adj. p-value <0.05, **Supp. Table 5**).

Deiter’s cells (DCs) were severely reduced in *Jag1*^*Ndr/Ndr*^ mice (**Fig3f-g)**, with a total of 30 cells compared to 126 cells in the *Jag1*^*+/+*^ dataset **(Supp. Table 8)**. Differential gene expression analysis identified 1 up- and 42-downregulated genes in *Jag1*^*Ndr/Ndr*^ DCs versus *Jag1*^*+/+*^ DCs (**Fig3h, Supp. Table 5,7**). The Notch target genes *Hes5* and *Nrarp* were upregulated (p-value <0.05) in *Jag1*^*Ndr/Ndr*^ DCs, suggesting increased Notch activation in mature *Jag1*^*Ndr/Ndr*^ DCs (**Fig3h-i, Supp. Table5**). There was no difference in DC signature genes (**Supp Table 7**), suggesting remnant *Jag1*^*Ndr/Ndr*^ DCs are correctly specified but exhibit elevated Notch activity at P5.

The reduction in lateral SCs, and lack of HeCs, could be a consequence of disrupted prosensory domain establishment in the absence of *Jag1*. At E14.5, *Jag1*^*Ndr/Ndr*^ mice exhibited similar SOX2^+^ and JAG1^+^ domains as *Jag1*^*+/+*^ mice (**Supp. Fig 2a-b**) but smaller lateral SOX2^+^JAG1^-^ lateral prosensory domains, and reduced expression of the Jag1-dependent prosensory domain marker CDKN1B (**Supp Fig 2a-b)**. Corroborating an overall reduction in Notch target gene expression during prosensory domain establishment, RNAscope demonstrated reduced expression of the Notch target genes *Heyl, Hey1, Hey2*, and *Hes1* in this domain (**Supp. Fig 2c-d**). In sum, a smaller initial *Jag1*^*Ndr/Ndr*^ lateral prosensory domain may limit development of lateral DCs and HeCs, but remnant medial *Jag1*^*Ndr/Ndr*^ DCs exhibit elevated expression of canonical Notch target genes.

### scRNAseq analysis demonstrates increased Notch activation in *Jag1*^*Ndr/Ndr*^ lateral SCs

As *Jag1*^*Ndr/Ndr*^ DCs exhibited elevated *Hes5* and *Nrarp* expression, we next investigated Notch component and target gene expression in *Jag1*^*Ndr/Ndr*^ and *Jag1*^*+/+*^ SCs and HCs. SCs expressed both Notch receptors and ligands, including *Notch1* and *Jag1* (**Fig3i**). Notch receptors were not detected in HCs, but *Jag2* was expressed in both *Jag1*^*Ndr/Ndr*^ and *Jag1*^*+/+*^ HCs (**Fig3i, Supp. Fig3**) and was significantly upregulated in *Jag1*^*Ndr/Ndr*^ OHCs (**Supp. Table5**), suggesting *Jag1* represses *Jag2* in OHCs.

To further study the Notch activation status of SCs, we cross-referenced differentially expressed genes in lateral SCs (IPCs, OPCs and DCs, avg. log2 FC >0.25 or <-0.25, and p-value <0.05) with genes previously reported to contain inducible binding sites for the Notch transcription factor Rbpj^35^. Of 258 reported target genes^35^, 27 were, in addition to *Jag1, Notch1* and *Nrarp*, differentially expressed in *Jag1*^*Ndr/Ndr*^ lateral SCs, of which most (20) were upregulated in lateral SCs, corroborating increased Notch activation in lateral SCs (**Fig 3I, Supp. Fig 3)**. In contrast to lateral SCs, HeCs do not express Jag1, but conditional deletion of *Jag1* results in the loss of HeCs^10^. Although the statistical power is limited due to the low HeC number in *Jag1*^*Ndr/Ndr*^ mice, the few remaining *Jag1*^*Ndr/Ndr*^ HeCs downregulated the Notch target gene *Nrarp* (**Fig3i**), suggesting lateral SCs (OPC, DCs) and HeCs do not respond similarly to *Jag1* insufficiency. In line with this, most of the Notch-inducible genes were upregulated in lateral SCs, but downregulated in HeCs, (**Fig3i**). In sum, we observed an overall upregulated Notch activation signature in lateral SCs, indicating a role for Jag1 in repressing Notch activity in lateral SCs.

### scRNAseq analysis of *Jag1*^*Ndr/Ndr*^ HCs reveals a role for Jag1 in OHC development and function

Finally, to determine whether Jag1 controls IHC or OHC specification, we analyzed *Jag1*^*Ndr/Ndr*^ and *Jag1*^*+/+*^ IHC and OHC transcriptomes. IHC transcriptomes were similar between *Jag1*^*Ndr/Ndr*^ and *Jag1*^*+/+*^ animals, with only five significantly dysregulated genes (**Supp. Table5**). Cross-referencing an extended list of *Jag1*^*Ndr/Ndr*^ IHC putatively dysregulated genes (322 genes, p-value <0.05) with the curated IHC signature (**Supp. Table 7**), revealed that only 4 of 322 genes (1.2%) were IHC-enriched. Therefore, even in presence of a medial border defect resulting in supernumerary IHCs and IPhCs (**Fig 3a-b, 4b-c**), the reduced Jag1 signalling does not significantly perturb IHC identity.

**Figure 4:**
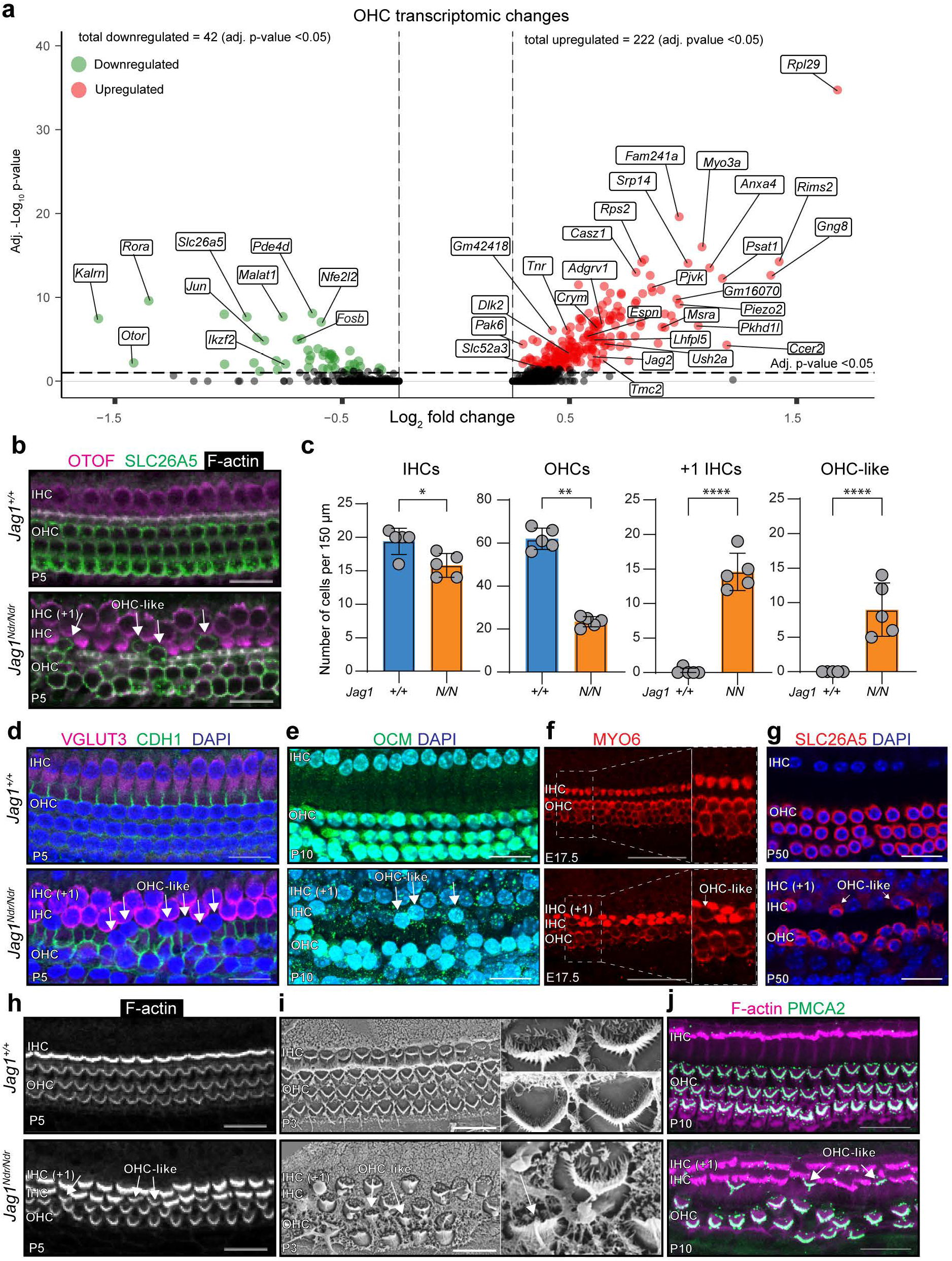
*Jag1*^*Ndr/Ndr*^ OHCs exhibit major transcriptomic dysregulation and HC patterning defects. **a)** Volcano plot depicting down-regulated (left, green) and up-regulated genes (right, red) for *Jag1*^*Ndr/Ndr*^ OHCs (adj. p-value <0.05). **b)** HC phenotype, showing IHCs (OTOF, magenta) and OHCs (SLC26A5, green) and PC protrusions (F-actin, white), showing a duplication of IHCs, fewer lateral OHCs, and the presence of SLC26A5-positive cells outside of the lateral OHC domain, referred to as OHC-like cells (white arrows). **c)** Quantification of IHCs, OHCs, supernumerary IHCs (IHC +1) and OHC-like cells. 14.6±2.7 +1 IHCs per 150 µm in *Jag1*^*Ndr/Ndr*^ compared to 0.2±0.5 in *Jag1*^*+/+*^, p-value <0.0001, 23.4±2.4 total OHCs per 150 µm in *Jag1*^*Ndr/Ndr*^ compared to 62.0±5.0 in *Jag1*^*+/+*^, p-value <0.0001, mean ± SD, 9.0±1.7 OHC-like cells per 150 µm in *Jag1*^*Ndr/Ndr*^ compared to 0.0±0.0 in *Jag1*^*+/+*^, p-value <0.001, mean ± SD. **d)** Phenotypic characterization of HC in *Jag1*^*Ndr/Ndr*^ mice (lower panels) at various stages of development. showing IHCs (VGLUT3, magenta), and E-cadherin (CDH1, green), indicating absent IHCs marker expression in OHC-like cells (white arrow), and the presence of OHC-like cells outside of the E-cadherin positive domain; **e)** Oncomodulin (OCM, green) expression in OHC-like cells. **f)** E17.5 cochleae stained for HCs (MYO6, red), demonstrating the presence of OHC-like cells in the inner/medial compartment, based on morphological characteristics. **g)** Panel showing OHCs (SLC26A5, red) at P50 (adult mice), with OHC-like cells in *Jag1*^*Ndr/Ndr*^ mice. **h**) Assessment of stereocilia (F-actin, white), using a different focal plane from the same cochleae as in (**b)**, demonstrating that *Jag1*^*Ndr/Ndr*^ OHC-like cells display V-shaped stereocilia and similar F-actin intensity similar to OHCs. **i)** Scanning electron microcopy (SEM) images of stereocilia, confirming OHC-like cell stereocilia resemble OHC stereocilia. **j)** OHC stereocilia-specific PMCA2 staining (green), demonstrating PMCA2-positive stereocilia outside of the lateral compartment in *Jag1*^*Ndr/Ndr*^ mice. n=5 or more per genotype; data are mean with standard deviation; scale bar represents 20 um; *p-value <0.05; **p-value <0.01; ***p-value <0.001, ****p-value <0.0001, unpaired t-test.

In contrast to IHCs, *Jag1*^*Ndr/Ndr*^ OHCs exhibited highly dysregulated gene expression, with 222 up- and 42 down-regulated genes (**Fig 4a**). Cross-referencing *Jag1*^*Ndr/Ndr*^ OHC dysregulated genes (264 genes, adj. p-value <0.05) with the curated OHC signature (**Supp. Table 7**), revealed that 40 of the dysregulated genes (15.2%) were OHC-enriched. In addition to *Jag2*, Notch pathway components *Dlk2* and *Hes6* were upregulated, as well as genes associated with inner ear and stereocilia development **(Fig 4a)**. Downregulated genes included mature OHC markers *Ikzf2* and *Slc26a5* (**Fig 4a**). Subsetting and renormalizing the HC subpopulation corroborated mature OHC marker downregulation, including *Ocm*, and upregulation of immature OHC makers such as *Bcl11b* and *Insm1 (***Supp. Fig 4a-b, Supp Table 9**). In sum, *Jag1*^*Ndr/Ndr*^ showed major transcriptomic dysregulation in OHCs, indicating a role for Jag1 in OHC gene regulation and development.

### *Jag1*^*Ndr/Ndr*^ mice exhibit OHCs in the IHC and PC compartments

Given the major transcriptomic changes in *Jag1*^*Ndr/Ndr*^ OHCs, we examined HC patterning in greater detail. While *Jag1*^*+/+*^ cochleae displayed the conserved one row of IHCs and three rows of OHCs, *Jag1*^*Ndr/Ndr*^ mice exhibited a complete second row of supernumerary IHCs (+1 IHC) (**Fig4b-d**), and the number of OHCs was greatly reduced, with an absent third OHC row **(Fig 4b-c)**. *Jag1*^*+/Ndr*^ mice exhibited an intermediate phenotype with the occasional supernumerary IHC, and occasional missing OHCs in the third OHC row (**Supp Fig 5a**), suggesting a dose-dependent IHC and OHC phenotype. Interestingly, *Jag1*^*Ndr/Ndr*^ mice exhibited sporadic ectopic expression of the OHC marker SLC26A5^+^ (Prestin) in HCs in the IHC and PC compartments (**Fig 4b, white arrows**). As these cells exhibited OHC marker expression, but were found outside of the OHC region, we refer to these ectopic cells as “OHC-like” cells. OHC-like cells were exclusively observed in *Jag1*^*Ndr/Ndr*^ mice but never in *Jag1*^*+/+*^ or *Jag1*^*+/Ndr*^ animals (**Fig 4c, Supp. Fig 5a-c**).

To further characterize OHC-like cells, we assessed their expression of an array of IHC- and OHC-specific markers. OHC-like cells were negative for the IHC markers Otoferlin (OTOF) and Vesicular glutamate transporter 3 (VGLUT3), as well as the outer compartment marker E-cadherin (CDH1) (**Fig 4d**) but expressed the OHC marker Oncomodulin (OCM) (**Fig 4e**). OHC-like cells were identifiable, based on morphological characteristics, as early as embryonic day (E) 17.5 (**Fig 4f, Supp. Fig 5b)**) and were still present after complete differentiation and maturation of the cochlea at P50 (**Fig 4g**). We next asked whether OHC-like cells would separate out as an additional cell population in the scRNAseq dataset. Renormalising and unsupervised reclustering of the HC dataset did reveal an OHC cell population exclusive to *Jag1*^*Ndr/Ndr*^ mice, with high expression of *Sez6l* and *Bmp2 (***Supp. Fig 6)**. However, RNAscope validation of *Sez6l* and *Bmp2* RNA expression demonstrated expression of these genes in PCs, rather than OHCs for both *Jag1*^*Ndr/Ndr*^ and *Jag1*^*+/+*^ animals (**Supp. Fig 6**). Given the location of some of the OHC-like cells in the PC compartment, they may be subject to a greater risk of contamination with PCs detritus during dissection and dissociation. Removing a PC gene signature (80 genes found in *Jag1*^*Ndr/Ndr*^ OHCs and in PCs of any genotype, **Supp. Table 9**), eliminated the separation of the *Jag1*^*Ndr/Ndr*^ OHC subpopulation, suggesting that the separation of this population could be driven by PC contamination, but also indicating that the OHC population of the *Jag1*^*Ndr/Ndr*^ dataset likely contains both (bridging) OHC-like cells and de facto OHCs, which are indistinguishable in this dataset in the absence of the PC signature (**Supp. Fig 6, Supp. Table 9**). The upregulation of immature OHC gene expression remained after PC signature removal, but pseudotime analysis indicated no statistically significant difference in pseudotime between *Jag1*^*Ndr/Ndr*^ and *Jag1*^*+/+*^ OHCs (**Supp. Fig 7)**, indicating that the developmental delay of *Jag1*^*Ndr/Ndr*^ OHCs, if any, is modest.

To further ascertain whether the ectopic OHC-like cells recapitulate the morphological and functional characteristics of OHCs, we next investigated morphology and molecular characteristics of stereocilia. All *Jag1*^*Ndr/Ndr*^ HCs were able to take up the FM1-43 dye, indicating that transduction channels in *Jag1*^*Ndr/Ndr*^ stereocilia are functional (**Supp. Fig 8)**. Moreover, OHC-like cells’ stereocilia had a V-shaped arrangement as well as low intensity F-actin staining, resembling *Jag1*^*+/+*^ OHCs (**Fig 4h**). Scanning Electron Microscopy (SEM) demonstrated a sparse arrangement of HCs in *Jag1*^*Ndr/Ndr*^ mice, with OHC-like cells exhibiting stereocilia bundle arrangement and characteristics of OHCs (**Fig 4i**). Finally, OHC-like cells’ stereocilia expressed the OHC stereocilia marker plasma membrane calcium pump Ca2^+^-ATPase 2 (PMCA2)^36^ (**Fig 4j**). Altogether, stereocilia on OHC-like cells recapitulate hallmarks of wildtype OHC stereocilia. In conclusion, ectopic *Jag1*^*Ndr/Ndr*^ OHC-like cells express markers and morphology of bona fide OHCs. Furthermore, a *Jag1*^*Ndr/Ndr*^ OHC sub-population, identified by a PC signature, clusters out upon sub-setting and re-clustering of the HC population.

### OHC-like cells exhibit a location-dependent expression of *Tbx2*

Factors determining an IHC vs OHC fate have recently been identified^37–43^, but it is not yet known whether Notch signalling controls expression of these genes in HC subtypes. We therefore investigated recently identified factors specifying these cell fates, focusing on the IHC determinant *Tbx2*^40^. We predicted that *Jag1*^*Ndr/Ndr*^ OHC-like cells should not express *Tbx2*, as its overexpression is sufficient to induce an IHC fate in OHCs, and ablation of *Tbx2* induces trans differentiation of IHCs to OHCs^40^. However, *Tbx2* was specifically expressed by the OHC-like cells found between IHCs (iOHC-like), but not by OHC-like cells bridging the IHC and OHC compartment, between PCs (bOHC-like) (**Fig 5**). In the presence of both *Tbx2* and *Slc26a5, Jag1*^*Ndr/Ndr*^ HCs are OHC-like, demonstrating that the expression of *Tbx2*, at least at the levels expressed in OHC-like cells, is not sufficient to consolidate an IHC fate in the *Jag1*^*Ndr/Ndr*^ cochlea.

**Figure 5:**
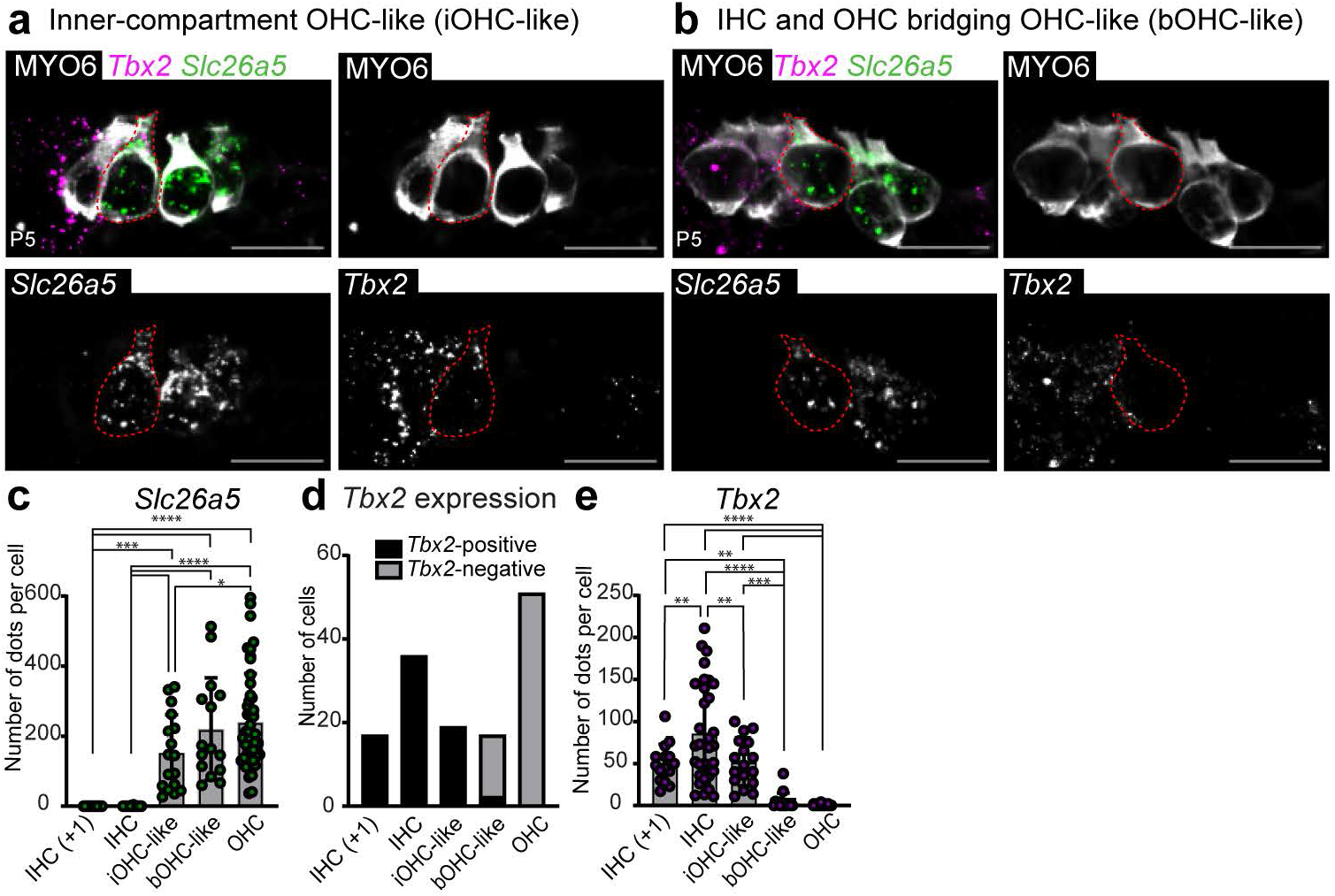
*Jag1*^*Ndr/Ndr*^ OHC-like cells exhibit location-dependent expression of the IHC fate determinant *Tbx2*. **a-b**) RNAscope for mRNA expression of the IHC regulator *Tbx2* (magenta) and OHC marker *Slc26a5* (green) in inner **(a)** and bridging OHC-like cells **(b)**, with HC counterstain (MYO6, white), demonstrating *Tbx2* and *Slc26a5* positivity in inner OHC-like cells and no *Tbx2* expression in bridging OHC-like cells (red encircled cells). **c)** Quantification of *Slc26a5* puncta in *Jag1*^*Ndr/Ndr*^ HC subtypes, 0.0±0.0 dots per IHC, 0.6±1.1 dots per +1 IHC, 153.2±108.3 dots per iOHC-like cell, 220.7±146.3 dots per bOHC-like cell and 240.4±138.2 dots per OHC, mean ± SD **d)** Quantification of *Tbx2* positivity/negativity (in binary terms) per HC subtype. **e)** Quantification of *Tbx2* mRNA puncta in *Jag1*^*Ndr/Ndr*^ HC subtypes, 85±58 dots per IHC, 50±22 dots per +1 IHC, 51±28 dots per iOHC-like cell, 4±10 dots per bOHC-like cell and 0.47±0.94 dots per OHC, mean ± SD, n= 5 or more per genotype; data are mean with standard deviation; scale bar represents 20 µm; *p-value <0.05; **p-value <0.01; ***p-value <0.001, ****p-value <0.0001, one-way ANOVA with Bonferroni correction.

In sum, *Jag1*^*Ndr/Ndr*^ mice exhibit atypical OHC-like cells in the IHC and PC compartments, in addition to supernumerary IHCs and a reduction in OHCs. OHC-like cells demonstrated a position-dependent expression of the IHC fate-determinant *Tbx2*, indicating that defective Jag1 signalling renders some HCs insensitive to the IHC fate determinant *Tbx2*.

## Discussion

In this study we utilized the Nodder mouse model of Alagille syndrome, to address how *Jag1* insufficiency affects individual SC and HC subtypes at single cell resolution. *Jag1*^*Ndr/Ndr*^ mice exhibited defects in prosensory domain size and Notch activation, medial boundary formation and lateral compartment defects. Importantly, OHC-like cells were present in the IHC or PC compartments and were insensitive to the IHC-determinant *Tbx2*, shedding new light on the potency of *Tbx2* in IHC fate determination in the context of defective Notch signaling. Together, our data demonstrates that *Jag1*^*Ndr/Ndr*^ mice provide a model hearing loss in ALGS and provide high-resolution insights into medial and lateral compartment development in the *Jag1*-compromised disease state.

### Boundary formation and lateral inhibition

Notch signalling establishes a boundary between the organ of Corti and Kolliker’s organ^44^, and patterns HCs and SCs via lateral inhibition. Partial reduction in Notch signalling results in a medial boundary defect with a proportional duplication of both a SC and a HC, while complete abrogation of Notch signaling results in both a boundary cell defect and profound lateral inhibition defects with a large excess of IHCs and no IPhCs^44^. Nodder mice exhibited a dose-dependent medial boundary defect that included a slight increase in IHCs in *Jag1*^*+/Ndr*^ mice (**Supp Fig 5**), and a full duplication of both IHCs and IPhCs in *Jag1*^*Ndr/Ndr*^ mice (**Fig 3,4**). The duplication and persistence of IPhCs show that lateral inhibition (by Dll1 or Jag2) is not lost in the *Jag1*^*Ndr/Ndr*^ medial domain. Similarly, in the lateral domain, whose size is likely restricted due to hampered prosensory domain establishment (**Supp Fig 2**), the reduction in OHCs was accompanied by a proportional reduction in the associated DC population, again suggesting preserved lateral inhibition. Interestingly, Notch signalling is specifically required for DC development and survival^19^, in which case DCs could be expected to be more severely reduced than OHCs in *Jag1*^*Ndr/Ndr*^ mice. Instead, *Jag1*^*Ndr/Ndr*^ DCs were reduced in proportion to the OHC reduction and exhibited a NOTCH-ON signature, with upregulation of multiple Notch target genes (**Fig 3h,i**). The increased Notch activation in *Jag1*^*Ndr/Ndr*^ SCs can be explained by two modes of Notch activation: 1) trans-activation from increased expression of Jag2 in *Jag1*^*Ndr/Ndr*^ OHCs (**Fig4a**) or 2) failure of JAG1^NDR^ to mediate Notch *cis*-inhibition within SCs. Although *cis-*inhibition is not thought to play a role during cochlear development, and research has been limited to Dll1^8,45^, it has been suggested that medial SCs can experience cis-inhibition^44^. The JAG1^NDR^ missense mutant is expressed *in vivo*, and traffics normally^46^, but does not bind or activate NOTCH1^14^. Our data does not allow us to conclusively determine whether Jag1 *cis*-inhibition or Jag2 *trans*-activation mediates the observed Notch activation in lateral *Jag1*^*Ndr/Ndr*^ SCs but indicates that great care must be taken to consider the levels of all Notch receptors and ligands in all cochlear cell types in these types of analyses. Experiments reducing *Jag2* levels in *Jag1*^*Ndr/Ndr*^ OHCs, by reducing *Jag2* gene dosage in *Jag1*^*Ndr/Ndr*^ mice, or by using JAG2 blocking antibodies, could theoretically address whether *Jag1*^*Ndr/Ndr*^ SC Notch activation is dependent on *Jag2* upregulation in OHCs, but would be technically challenging due to the required specific mixed genetic background of *Jag1*^*+/Ndr*^mice^14^, their vascular frailty ^47^, and their high mortality^14,47^

### Origin of OHC-like cells

OHC and IHC specification in the lateral and medial compartments, respectively, is a tightly regulated process, for which only a few perturbations have recently been described^39–43,48^. The presence of OHC-like stereocilia on both iOHC-like cells and bOHC-like cells, including the V-shape bundle arrangement, F-actin intensity levels and PMCA2 positivity argue against a direct IHC-to-OHC-like cell conversion (**Fig 4h-j**). Conditional deletion of *Tbx2* induces neonatal transdifferentiation of IHCs into OHCs, expressing several OHC molecular markers but lacking OHC stereocilia features. Instead, the transdifferentiated cells maintained stereocilia that resemble the overall appearance of IHC stereocilia ^40^. HC stereocilia develop during a limited time window during a late embryonic and early postnatal period and do not alter stereocilia size and bundle arrangement once stereocilia are formed, and are not replaced once lost^49,50^, indicating that *Jag1*^*Ndr/Ndr*^ OHC-like cells most likely were born as OHCs rather than converted from IHCs into OHC-like cells. Although our experiments do not directly address the developmental origin of the *Jag1*^*Ndr/Ndr*^ OHC-like cells, published data, their anatomical location, and transcriptomic data suggest that bOHC-like cells might originate from PCs or their precursors. Ectopic HCs in the PC region are present at the position of missing PCs in *Hey1, Hey2* double mutant mice, suggesting a late PC-to-HC conversion^51^. *Jag1*^*Ndr/Ndr*^ OHC-like cells appear before IPC and OPC differentiation, and PC differentiation is delayed in *Jag1*^*Ndr/Ndr*^ mice. Based on the position of the bOHC-like cells, the PC precursor population could be a source of cells that differentiate into OHC-like cells. However, SCs that convert into HCs typically retain at least a partial SC gene expression signature, including low levels of SOX2^52^, which we did not observe at various stages using SOX2 immunostaining, nor in any of the scRNAseq OHC populations. Nonetheless, OHC subsetting revealed a *Jag1*^*Ndr/Ndr*^-specific OHC population expressing a PC signature (**Supp Fig 6**), which could reflect either contamination of *Jag1*^*Ndr/Ndr*^ cells specifically due to their anatomical location or a divergent developmental origin. Therefore, OHC-like *Jag1*^*Ndr/Ndr*^ cells are more likely to have originated from PCs, or a PC precursor pool, than from IHCs, and future studies utilizing for example time-lapse imaging of developing *Jag1*^*Ndr/Ndr*^ cochlear explants or lineage tracing could help resolve whether iOHCs and bOHCs share a developmental origin, and whether these arise from IHCs, PCs, or other cells.

### Potency of cell fate determinants

Recently, the identification of IHC-versus OHC-specific factors including *Tbx2*^39–41^, *Insm1*^42,43^ and *Ikzf2*^48^ have significantly contributed to understanding HC divergence into IHC and OHCs and highlighted neonatal plasticity between the two cell types. Although an interplay between Notch and these factors have been suggested in other tissues^53,54^, it is not reported whether Notch signaling regulates these factors in the cochlea. Previous work has demonstrated that *Tbx2* is epistatic to *Insm1*^42^: in the absence of both *Tbx2* and *Insm1*, only OHCs develop, demonstrating that *Tbx2* is required for normal IHC differentiation and trans differentiation of Insm1-deficient OHCs into IHCs. Ectopic *Tbx2* expression in P0 cochlear explant OHCs represses OHC features, including Prestin expression, and induces IHC features, including VGLUT3 expression^40^. However, *Jag1*^*Ndr/Ndr*^ mice exhibited *Tbx2*^+^ iOHC-like cells, indicating that *Tbx2* is not sufficient to induce an IHC fate in iOHC-like cells in the absence of functional Jag1. Importantly, *Tbx2* is an organ of Corti inner compartment patterning factor^41^, which may explain its expression in *Jag1*^*Ndr/Ndr*^ OHC-like cells in the IHC compartment. It is possible that Tbx2 dosage is important for consolidating the IHC fate: *Tbx2* levels were lower in iOHC-like cells compared to IHCs, which might permit the OHC-like phenotype (**Fig5e**). However, the supernumerary IHCs (IHC+1) expressed similar levels of *Tbx2* as iOHC-like cells but developed as IHCs rather than OHCs (**Fig5e**), arguing against Tbx2 gene dosage as the sole determining factor in *Jag1*^*Ndr/Ndr*^ OHC-like phenotype. More recently, it has been shown that *Tbx2* overexpression alone is insufficient to induce an IHC fate in medial SCs^39^, indicating that further studies are required to understand the potency of Tbx2 in driving an IHC fate.

#### Hearing and balance in Alagille syndrome

Both conductive and sensorineural hearing loss are reported in patients with Alagille syndrome^5^, as well as vestibular defects^6^. Similarly, *Jag1*-deficiency results in hearing loss in mice ^9–12^ (**Supp Table2**), and *Jag1* missense mutant mice are nicknamed for overt balance defects, including *Headturner*^18^, *Ozzy*^11^ and *Slalom*^17^. ABR measurements indicated severe hearing loss in *Jag1*^*Ndr/Ndr*^ mice, as well as reduced cochlear amplification by DPOAE. *Jag1*^*Ndr/Ndr*^ IHC transcriptomes demonstrated very few dysregulated genes. The ABR-related hearing defects in *Jag1*^*Ndr/Ndr*^ mice are thus unlikely to be caused by IHC defects, as reported for mice with SC-specific Jag1 deletion^9^. Instead, reduced cochlear amplification, middle ear bone malformation, and a reduction in OHCs accompanied by pronounced transcriptomic changes in OHCs, could together explain the severe hearing loss in *Jag1*^*Ndr/Ndr*^ mice.

Phenotype presentation in ALGS is generally diverse, and there is no reported genotype-phenotype correlation^14,47,55,56^. While features of ALGS such as intrahepatic bile duct paucity and vascular defects similarly exhibit variability in *Jag1*^*Ndr/Ndr*^ mice^14,47,56^, the inner ear defects were remarkably consistent within *Jag1*^*Ndr/Ndr*^ and *Jag1*^*Ndr/+*^ mice. The inner ear may thus provide a model that is less sensitive to environmental modifiers, or genetic modifiers in the mixed C3H/C57bl6 background that the Nodder colony must be maintained in. *Jag1*^*Ndr/Ndr*^ mice thus present a relevant disease model in which to test potential therapeutics and demonstrate that a genotype-phenotype correlation (or dose-dependency) might be present in some organs, and absent in others.

#### Concluding remarks

In conclusion, this study reveals new functions for *Jag1* in cochlear cell patterning and cell identity, with single cell resolution. In addition to previously reported phenotypes, we identified and characterized a novel *Tbx2 and Slc26a5* double positive OHC-like cell in the medial compartment, shedding new light on the potency of *Tbx2* in IHC fate determination in the context of defective Notch signaling. The scRNA seq dataset provides an atlas of the Notch-dysregulated cochlea, and reveals major gene dysregulation in OHCs, as well as a novel role for Jag1 regulating Notch activation in lateral SCs. By characterizing patterning defects, as well as transcriptomic changes in *Jag1*^*Ndr/Ndr*^ mice, these data lay a foundation for future investigation of the interaction between Notch signaling and sensory cell patterning and specification, as well as HC divergence and regeneration, and demonstrate that *Jag1*^*Ndr/Ndr*^ mice provide a model for ALGS with hearing and balance defects.

## Materials and Methods

### Animals

*Jag1*^*+/Ndr*^mice originate from the ENU induced mutation program as described previously^46^. Mice were maintained on a C3H/C57BL6 mixed background as described^14^. F2 generation mice were used for this study. All experiments were performed in accordance with EU rules and regulations as approved by the Swedish Board of Agriculture (Jordbruksverket) under ethical permit numbers N5253/2019 and N2987/2020. To obtain timed embryonic and postnatal animals, animals were mated overnight and noon of the day of vaginal plug was considered embryonic day (E) 0.5. Animals were maintained under standard day/night cycles and provided with food and water ad libitum and housed in enriched individually ventilated cages (wood, paper embedding and cardboard tunnels), with a maximal number of six animals per cage. Each mouse used was genotyped by Transnetyx (Cordova, TN).

### Open Field Test – Circling analysis

Male and female *Jag1*^*+/+*^ and *Jag1*^*Ndr/Ndr*^ mice (6-8 mice per sex and genotype) at 3-12 months were used for the Open Field Test. Mice were gently placed in a plexiglass openfield box and filmed with a TSE VideoMot2 system. Distance travelled in pixels was converted to actual distance using a scale measure. Each mouse was filmed for 30 minutes, and the box was cleaned with water and dried in between each mouse. The experimenters were two different female scientists, who sat still during the experiments.

### Auditory measurements

All auditory measurements were performed using Tucker-Davis Technologies System III hardware and software, and stimuli were generated using BioSigRZ software connected to a signal processor (RZ6). For DPOAEs, sound from two independently driven MF1 speakers was merged into a custom acoustic coupler for closed field stimulation. For ABR, the stimulus output was through an open field speaker (MF1). A Brüel and Kjær ¼-inch microphone and a conditioning pre-amplifier (4939 A 011 and 2690 A 0S1) were used to calibrate the stimulus. Speakers were calibrated one at a time using a frequency sweep (4 to 32 kHz). The output was corrected to produce a flat spectrum at 90-dB SPL (open field speaker, ABR) and 80-dB SPL (closed field speakers, DPOAEs). Calibration of speakers was performed daily prior to auditory measurements. Sex-matched animals from different genotypes in age range P45-P60 were anesthetized using a mixture of ketamine [ketaminol (50 mg/ml), Intervet, 511485] and xylazine [Rompunvet (20 mg/ml), Bayer, KPOCRD5] (100 and 10 mg/kg body weight, respectively) by intraperitoneal injection and placed in a custom-made acoustic chamber. Body temperature was maintained at 36.5°C using a heating pad (Homeothermic Monitoring System 55-7020, Harvard Apparatus) and eye gel was applied to prevent ocular dehydration during anesthesia. DPOAEs were recording prior to ABR measurement, by placing an acoustic coupler into the ear canal. A microphone (EK 23103, Knowles) was inserted into the acoustic coupler and connected to a pre-amplifier (ER-10B+, Etymotic Research) and a processor (200-kHz sample rate) to measure sound intensity in the ear canal. Each speaker played one of two primary tones (f1 and f2) and swept in 5-dB steps from 80- to 10-dB SPL (for f2). The 2f1 − f2 distortion product was measured with f2 = 8, 12, 16, 24, and 32 kHz, f2/f1 = 1.25, and stimulus intensity L1 = L2 + 10-dB SPL. Following DPOAE, ABRs were measured by subdermal placement of stainless-steel needle electrodes at the head vertex (positive), under the right ear pinna (negative) and above the right leg (ground). ABRs were evoked by tone bursts (0.5 ms rise per fall time, 5-ms duration) of 8, 12, 16, 24, and 32 kHz presented 21 times per second. Signals were collected via a low-impedance head stage (RA4LI) connected to a pre-amplifier (RA4PA) and digitally sampled (200-kHz sample rate). Responses to 1000 bursts were bandpass filtered at 0.3 to 3 kHz using BioSigRZ and averaged at each intensity. For each frequency, sound intensity decreased from 90-dB SPL in 5-dB steps. Repeated injection of anesthesia, at ¼ of initial dose, was occasionally given during the ABR measurement to ensure stable anesthesia throughout the measurement. ABR thresholds were determined by the lowest sound stimulus for which a reproducible ABR waveform was identified. DPOAE thresholds were determined as the lowest sound stimulus by which a DPOAE amplitude reaches 5 dB above noise.

### Tissue dissociation and library preparation for single cell RNA sequencing

Pups were decapitated at postnatal day 5 and cochleae were collected. The sensory epithelium and surrounding mesenchyme were dissected out. Two cochlea from each pup were incubated for 15 minutes in 3ml DMEM:F12 with 100 μl Thermolysin (Sigma, 5 mg/ml) and 5 μl DNase (1 mg/ml Stemcell Technologies) at 37°C. Cells were dissociated in pre-activated Papain (20 U/mL, Worthington Papain Dissociation Kit, Cat No. LK003150, activated by placing in 37°C for 30 minutes with open lid) for a total of 15-20 minutes with trituration every 5 minutes using 200 μl low bind tips (Sartorius, 790101F). Papain was inactivated using an Ovomucoid inhibitor (Worthington Papain Dissociation Kit) and dissociated cells were passed through a 20 μm cell strainer, spun down at 300x g and resuspended in 500 μl 0.4 mg/mL BSA (Ambion Ultrapure BSA, AM2616). Propidium Iodide (ThermoFisher, P1304MP) was added to the cell suspension (final concentration 50 μg/ml) and cells were sorted using a BD FACS Aria cell sorter at 4°C and collected in 0.4 mg/mL BSA in DNA LowBind tubes (Eppendorf, 0030108523). Cells were pelleted at 300 g for 5 minutes at 4°C and manually counted using a Bürker Chamber. Following single cell dissociation, single cells were captured and partitioned into droplets using a 10X Chromium controller, and libraries were prepared using Chromium Next GEM Single Cell 3′ Reagent Kits v3.1 according to the manufacturer’s instructions. cDNA and post library quality controls were performed using an Agilent Bioanalyser and DNA concentrations were determined using Qubit dsDNA (ThermoFisher). Libraries were sequenced on an Illuminia NextSeq550 System, using NextSeq 500/550 High Output Kit v2.5, 150 cycles flow cells (Illumina, 20024907). Library preparations of one sample per genotype were run on a single flow cell, using the following read set-up: read 1: 28 cycles; i7: 8 cycles, i5: not used, single index only; read 2: 130 cycles.

### Single cell RNAseq data processing

FASTQ files were aligned to a reference genome GRCm38 using CellRanger 5.0. CellRanger h5 output files were imported and converted into Seurat Objects using Seurat V4. Quality control was performed for each dataset, excluding cells with the following features: >10% mitochondrial genes, > 50.000 RNA counts, < 2000 or > 10.000 genes or features. After initial quality control, doublets as indicated by scDbFinder package (version 1.14.0) were removed from each dataset. The expression data were then normalised using SCTransform V2, with regression out of mitochondrial genes. Principal components analysis (PCA) and Uniform Manifold Approximation and Projection (UMAP) was used as Non-linear dimensionality reduction technique for cell clustering. Epcam^+^ clusters were identified and subsetted for further analysis. Datasets were integrated using Seurat SCTransform V2 “IntegrateData” and dimensionality reduction settings were determined using “ElbowPlot” and examination of maker gene expression for different clusters. Subsetting of specific clusters (DC and PC) was performed to separate different cell types and were assigned according to the parental UMAP. A subset of DCs from the wildtype dataset was excluded based on high expression levels of periotic mesenchyme markers *Tbx18* and *Pou4f3* and was considered to be contamination (for full code availability see https://github.com/Emma-R-Andersson-Lab). Pseudotime analysis was performed using Monocle (version 1.3.1)

### Immunohistochemistry (wholemount preparations)

Embryos and pups were collected from timed pregnant females at various embryonic and postnatal timepoints, as indicated. Inner ears were dissected out from the skull and fixed in fresh 4% PFA for 1.5 hours at room temperature. Fixed inner ears were washed in PBS overnight at 4°C. Otic cup, Reisner’s membrane and the tectorial membrane were subsequently dissected off to expose the sensory epithelium of the cochlea. Cochleae were blocked in PBS containing 10% Normal Donkey Serum (Sigma, Cat. No D9663) and 0.3% TritonX (Sigma, Cat. No. T8787) for 1 hour at room temperature. Cochleae were incubated in primary antibody overnight at 4°C (for primary antibody details, please see **Supp. Table 10**). Cochleae were washed 3 times for 15 minutes in PBS and incubated in secondary antibodies in PBS for 1 hour at room temperature. After secondary antibody incubation, cochleae were washed 3 times for 15 minutes in PBS and the sensory epithelium was dissected off and mounted (Sigma-Aldrich, Cat. No. F6182) on a glass slide. Secondary antibodies were used at 1:500 Donkey anti-Rabbit Alexa Fluor 488 (Abcam, AB150073), Donkey anti-Goat Alexa Fluor 546 (ThermoFisher, A11056), Donkey anti-Goat Alexa Fluor 594 (Abcam, AB150132), Donkey anti-Sheep Alexa Fluor 555 (Abcam, 150178) and DAPI (Sigma, MBD0015) for nuclear counterstain.

### RNAscope (cryosections)

Embryos and pups were collected from timed pregnant females at various embryonic and postnatal timepoints, as indicated. Inner ears were dissected out from the skull and fixed in fresh 4% PFA overnight in the cold room. Cochleae were exposed to a sucrose gradient (5, 10, 15, 20, 25% sucrose, 30 minutes each) and incubated overnight in 30% sucrose at 4°C. Sections were embedded in O.C.T. medium and stored at -80°C. 12 µm tissue sections were made using a Epredia CryoStar NX70 Cryostat (blade temperature -20°C) NX70 cryostat. RNAscope hybridisation was performed according to the manufacturer’s instructions (ACDbio). Following RNAscope, sections were incubated with primary antibodies and processed as described in the previous section. For RNAscope probe details, please see **Supp. Table 10**.

### Image acquisition and quantification

Confocal images were acquired using LSM880 or LSM980 with Zeiss ZEN software. Acquisition parameters were selected to maximize contrast between structures of interest. The sharpest focal plane with the clearest structures was selected for analyses. HCs and SCs were manually counted using ImageJ software. For prosensory domain size measurements, Qupath 0.4.4 or Zeiss Zen software was used using “Polygon” or “Draw spline contour” tool, respectively. For quantification of RNAscope signal, images were acquired using the same imaging settings among samples, and a customized ImageJ script was used by adjusting images to a threshold followed by “Measure Particles”. The number of particles was subsequently multiplied by the average particle size and normalized by dividing by the size of the smallest particle per cochlea, as recommended by manufacturer’s technical note (TS-46-003).

### Scanning electron microscopy

Cochleae were dissected out and otic cup was removed. Tissue was fixed in fixation buffer (2.5% glutaraldehyde, 2% paraformaldehyde in 0.1 M phosphate buffer) for 1 hour at room temperature and stored in buffer at 4°C. Following fixation, the sensory epithelium was removed and trimmed for final preparation. Cochleae were dehydrated using a series of ethanol gradients (20-40-60-80-100% ethanol) and were critical point dried using liquid CO2. Dried samples were mounted on SEM mini-holders, sputter-coated with platinum, and imaged using a Zeiss Gemini Ultra 55 SEM.

### FM1-43 dye staining

Inner ears were dissected out and the otic cup was removed to expose the sensory epithelium. Fresh inner ears were incubated for 60 s in FM1-43 (Thermofisher, F35355) in PBS (final concentration 5 µM) and washed with 1x PBS and kept in the dark. After washing, cochleae were fixed in 4% PFA and processes as described in the immunohistochemistry section.

## Supporting information

Graphical abstract

## Acknowledgements

We acknowledge and thank the following funders: KI-NIH doctoral programme fund, Karolinska Institutet Career Grant (KI Consolidator to ERA), the Swedish Research Council (ERA), The Swedish Foundations’ Starting Grant, provided by The Ragnar Söderberg Foundation (ERA) and the Swedish Research Council (ERA). We thank Evangelia Tserga and Corstiaen Versteegh for assistance with auditory measurements, and Igor Baars for sequencing assistance. We thank Saida Hadjab and Susi Gralla for advice and help with the Open Field measurements. We thank Göran Månsson and Florian Salomons from the Karolinska BIC facility for advice on image acquisition, and the technical staff at KM- A/B animal facilities from Comparative Medicine for colony handling. We thank Tessa Sanders, Helen Maunsell, Braulio Peguero and Beatrice Mao for practical training on inner ear dissections and processing (SDH) and scientific discussions.

## Author contributions

Sandra de Haan: Conceptualization, experimentation, analysis, investigation and validation, intellectual input, original draft writing and revision. Agustin A. Corbat: Analysis, intellectual input, reviewing. Christopher R. Cederroth: Supervision, intellectual input, reviewing. Lisa G. Autrum: data collection. Simona Hankeova: data collection, reviewing. Elizabeth C. Driver: intellectual input and reviewing. Barbara Canlon: reviewing: Matthew W. Kelley: Supervision, conceptualization, analysis, intellectual input, Emma R. Andersson: Supervision, data collection, conceptualization, analysis, intellectual input, original draft writing and revision.

