## Supplement Table1 for "Jag1 represses Notch activation in lateral supporting cells and inhibits an outer hair cell fate in the medial compartment of the developing cochlea"

**Supplement Table 1:** Summary of Notch defective animal models reporting an inner ear phenotype, extended table of Brown et al Table1^1^

| **Receptors** | **Type of mutation** | **Phenotype** | **Ref** |
| --- | --- | --- | --- |
| ***Notch1*** | Inner ear specific knockout  (Pax2-cre) | Extra OHC row, correctly patterned | ^2^ |
| ***Notch1*** | Inner ear specific knockout  (Foxg1-cre) | 3-fold increase in IHCs and OHCs, disorganized | ^3^ |
| ***Notch1*** | SC specific knockout  (Sox2CreER x Notch1fl/fl) at P0/P1 | Loss of lateral SCs, and HC loss secondary to SCs loss. | ^4^ |
| **Ligands** | **Type of mutation** | **Phenotype** | **Ref** |
| ***Jag1*** | Headturner (*Htu*^+/-^)  ENU-induced missense mutation (G289D) | Ectopic IHCs, reduction OHCs | ^5^ |
| ***Jag1*** | Ozzy (*Ozzy*^+/-^)  ENU-induced missense mutation (W167R) | Ectopic IHCs, reduction OHCs | ^6^ |
| ***Jag1*** | Slalom (*Slm*^+/-^)  ENU-induced missense mutation (P269S) | Ectopic IHCs, reduction OHCs | ^7^ |
| ***Jag1*** | *Inner ear specific knockout*  *(Foxg1-cre)* from E8.75 | Duplication of IHCs, lack of OHCs in mid cochlea | ^8^ |
| ***Jag1*** | SC specific knockout  (*Sox2^CreERT2^/Jag1^loxP/loxP^)* | Loss of HeC | ^9^ |
|  | Inner ear specific knockout  (*Fgfr3^iCreERT2^/Jag1^loxP/loxP^*) | Loss of HeC | ^9^ |
| ***Jag1*** | SC specific knockout  (*Sox2^CreER^/Jag1^fl/fl^)* | Loss of some HeC | ^4^ |
| ***Jag2*** | Null mutant, heterozygous  *(Jag2*^+/DSL^ ) | Ectopic IHCs | ^10^ |
| ***Jag2*** | Null mutant, homozygous  *Jag2*^DSL /DSL^ | Duplication of IHCs, extra row of OHC, abnormal orientation of stereocilia on OHCs. | ^11^ |
| ***Dll1*** | Null mutant, heterozygous  *(DLL1*^Dll1ki/+^) | Ectopic IHCs and supernumerary OHCs | ^12^ |
| ***Dll1*** | Null mutant, homozygous  (*DLL1*^Dll1ki/LacZ^) | Ectopic IHCs and supernumerary OHCs | ^12^ |
| ***Dll1*** | Inner ear specific knockout  *(Foxg1-cre)* | Ectopic IHCs, supernumerary OHCs, and delayed cochlear growth | ^13^ |
| ***Dll1+ Jag2*** | Double null mutant, heterozygous  *(Jag2*^+ /DSL^ / *DLL1*^+/LacZ)^ | Duplication of IHCs, extra row of OHCs | ^12^ |
| ***Dll1+ Jag2*** | Double null mutant,  (*Jag2*^DSL /DSL^ / *DLL1*^+/LacZ^) | Duplication of IHCs, extra row of OHCs, disorganised | ^12^ |
| ***Dll1+ Jag2*** | Double null mutant,  (*Jag2*^DSL /DSL^ / *DLL1*^+/Dll1ki^) | Two to four rows of IHCs, four to six rows of OHCs, disorganised | ^12^ |
| ***Dll3*** | Null mutant  *(DLL3*^pu/pu^) | No phenotype | ^14^ |
| **Modulators** | **Type of mutation** | **Phenotype** | **Ref** |
| ***Lfng*** | Null mutant  *(Lfng^LacZ/LacZ^ )* | No phenotype | ^2^ |
| ***Mfng*** | Null mutant  *(Mfng* ^Tm1.1Cfg/tm1.1Cfg^) | No phenotype | ^15^ |
| ***Lfng + Mfng*** | Double null mutant  *(Lfng^LacZ/LacZ^/ Mfng* ^Tm1.1Cfg/tm1.1Cfg^) | Ectopic IHCs and Iphcs | ^15^ |
| ***Lfng + Jag2*** | Double null mutant  *Jag2*^ΔDSL /ΔDSL^ / *Lfng*^lacZ/lacZ^ | Supernumerary OHCs, and abnormal stereocilia on OHCs. Rescues Jag2 IHC phenotype | ^16^ |
| ***Pofut1*** | Null mutant  (Pofut1^-/-^) | Ectopic IHCs | ^15^ |
| ***Pofut1*** | Inner ear specific knockout  (Pax2-cre) | Ectopic IHCs and Iphcs, supernumerary OHCs | ^17^ |
| **Downstream targets** | **Type of mutation** | **Phenotype** | **Ref** |
| ***Hes1*** | Null mutant, heterozygous  (Hes1^+/-^ ) | Ectopic IHCs | ^18^ |
| ***Hes1*** | Null mutant, homozygous  (Hes1^-/-^ ) | Duplication IHCs | ^18^ |
| ***Hes5*** | Null mutatant, homozygous  (Hes5^-/-^ )  (null mutation) | Supernumerary OHCs | ^18^ |
| ***Hes5 + Hes1*** | Double null mutant  (Hes1^-/-^ / Hes5^+/-^ ) | Duplication of IHCS | ^18^ |
| ***Hes5 + Hes1*** | Double null mutant  (Hes1^+/-^ / Hes5^-/-^) | Ectopic IHCs and supernumerary OHCs | ^18^ |
| ***Hey2*** | Null mutant, homozygous  (Hey2^-/-^) | Supernumerary OHCs | ^19^ |
| ***Hey2 + Hes5*** | Double null mutant, homozygous  (Hey2^-/-^/ Hes5^-/-^) | Supernumerary OHCs | ^19^ |
| ***Hey2 + Hes5*** | Double null mutant, homozygous  (Hey2^-/-^/ Hes5^-/-^) | Supernumerary OHCs | ^19^ |
| ***Hey2*** | Null mutant, homozygous  (*Hey2^-/-^)* | No phenotype, but additional DAPT treatment results in loss of PC and DCs. | ^20^ |
| ***Hey2*** | Null mutant, homozygous  (*Hey2^-/-^)* | No phenotype | ^21^ |
| ***Hey2 + Hes1*** | Double null mutant  (*Hey2^+/-^/ Hes1^-/-^* ) | Ectopic IHCs and supernumerary OHCs | ^19^ |
| ***Hey1*** | Floxed knockout  (*Hey1^fl/fl^*) | No phenotype | ^21^ |
| ***Hey1 + Hey2*** | Double null mutant  *Hey1^-/-^/ Hey1^-/-^* | Ectopic HCs in the PC region | ^21^ |
| ***Co-activators*** | **Type of mutation** | **Phenotype** | **Ref** |
| ***dnMaml*** | Inner ear specific knockout  (Pax2-cre) | Ectopic IHCs and Iphcs | ^15^ |
| ***dnMaml*** | Inner ear specific knockout  (Pax2-cre) | Loss of DCs | ^22^ |
| ***Rbpj*** | Inner ear specific knockout  (Foxg1-cre) | Mice die before before pattern establishment | ^23^ |
| ***Rbpj*** | Inner ear specific knockout  (Pax2-cre) | Mice die before before pattern establishment | ^24^ |
| ***Rbpj*** | Inner ear specific knockout  (Fgfr3-iCreER; Rbpj^−/Δ^) | Loss of DCs | ^22^ |
