## Supplement Table2 for "Jag1 represses Notch activation in lateral supporting cells and inhibits an outer hair cell fate in the medial compartment of the developing cochlea"

**Supplementary Table2: Summary of hearing function and middle ear bone morphology in *Jag1* mutant mouse models**

| **Model** | **Hearing function** | **Middle ear bones** | **Ref** |
| --- | --- | --- | --- |
| Conditional knockout  *(Wnt1-Cre*; *Jag1^f/f^)* | Elevated hearing thresholds among all frequencies | Malformed (columnar) stapes and occasional ectopic processes on incus | ^1^ |
| Headturner (*Htu*^+/-^)  ENU-induced missense mutation (G289D) | Elevated hearing thresholds, not significant | Not assessed | ^2^ |
| Ozzy (*Ozzy*^+/-^)  ENU-induced missense mutation (W167R) | Slightly elevated hearing thresholds, most pronounced at middle frequencies | Not assessed | ^3^ |
| Slalom (*Slm*^+/-^)  ENU-induced missense mutation (P269S) | Not assessed | Not assessed | ^4^ |
| *Inner ear specific knockout*  *(Foxg1-cre)* from E8.75 | Not assessed | Not assessed | ^5^ |
| SC specific knockout  (*Sox2^CreERT2^/Jag1^loxP/loxP^)* | Not assessed | Not assessed | ^6^ |
| Inner ear specific knockout  (*Fgfr3^iCreERT2^/Jag1^loxP/loxP^*) | Elevated thresholds, deafness at lower frequency range | Not assessed | ^6^ |
| SC specific knockout  (*Sox2^CreER^/Jag1^fl/fl^)* | Deaf at all frequencies | Not assessed | ^7^ |
