## Supplement Table10 for "Jag1 represses Notch activation in lateral supporting cells and inhibits an outer hair cell fate in the medial compartment of the developing cochlea"

Primary antibodies

| **Antibody** | **Vendor** | **Cat. Number** | **Dilution** |
| --- | --- | --- | --- |
| Cd44 | BD Biosciences | 550538 | 1:500 |
| E-cadherin | BD Biosciences | 610182 | 1:500 |
| FABP7 | R&D | AF3166 | 1:1000 |
| Jag1 | Cell. Sig.28H8 | 2620 | 1:125 |
| Myosin6 | BioProteus | 25–679 | 1:1000 |
| NGFR | R&D | AF367 | 1:500 |
| Ocomodulin | Novus Biologicals | NBP2-14568 | 1:500 |
| P27kip1 | BD Biosciences | 610242 | 1:250 |
| PCMA2 | Invitrogen | PA1-915 | 1:1000 |
| Phalloidin | ThermoFisher | A22287 | 1:500 |
| Slc26a5 | Abcam | Ab242128 | 1:20.000 |
| Sox2 | R&D | AF2018 | 1:250 |
| Vglut3 | Millipore | AB5421-I | 1:1000 |

RNAscope probes

| **Antibody** | **Vendor** | **Cat. Number** |
| --- | --- | --- |
| Hes1 | ACDbio | 417701-C1 |
| Hey1 | ACDbio | 319021-C3 |
| Hey2 | ACDbio | 404651-C2 |
| HeyL | ACDbio | 446881-C1 |
| Slc26a5 | ACDbio | 521321-C1 |
| Sez6l | ACDbio | 492631-C3 |
| Bmp2 | ACDbio | 406661-C2 |
| Tbx2 | ACDbio | 448991-C2 |
