## Supplement Legends for "Jag1 represses Notch activation in lateral supporting cells and inhibits an outer hair cell fate in the medial compartment of the developing cochlea"

**Supplementary Figure Legends**

**Supp. Fig 1: *Jag1^Ndr/Ndr^* mice show sporadic remnant HeCs and have delayed PC development. a)** *Jag1^+/+^* HeC phenotype showing HeCs (FABP7, green, also labels IPhCs), HCs (MYO7A, red) and nuclei (DAPI, blue) showing FABP7-positive cell rows in the IHC/IPhC and HeC positions. **b)** Representative image (left) of *Jag1^Ndr/Ndr^* HeC phenotype stained for HeCs (FABP7, green), HCs (MYO7A, red) and nuclei (DAPI, blue) showing the absence of FABP7-positive cell layers lateral to OHCs, in the HeC position, however, sporadic FABP7^+^ HeCs can be found (right). **c)** PC phenotype at P0 and E17.5, showing IPCs (NGFR, green) and OPCs (CD44, magenta) demonstrating widespread NGFR signal in P0 *Jag1^Ndr/Ndr^*  mice (left**),** resembling E17.5 *Jag1^+/+^* NGFR signal (right). **d**) At P10, the number of IPCs and OPCs is similar in *Jag1^Ndr/Ndr^* and *Jag1^+/+^* mice **c**) Quantification of PCs for P0 apical region, indicating an increase in NGFR positive cells, and a reduction of CD44 positive cells 40.2± 3.1 NGFR+ cells per 150 µm in *Jag1^Ndr/Ndr^* compared to 29.2±1.8 NGFR+ cells per 150 µm in *Jag1^+/+^,* p-value <0.001, mean ± SD and 1.2±1.1 Cd44+ cells per 150 µm in *Jag1^Ndr/Ndr^* compared to 23.8±2.8 CD44+ cells per 150 µm in *Jag1^+/+^,* p-value <0.001, mean ± SD. At P5, base region, the number of IPC and OPCs is similar for *Jag1^Ndr/Ndr^* and *Jag1^+/+^* mice 16.4±1.7 IPC cells per 150 µm in *Jag1^Ndr/Ndr^* compared to 19.2±2.3 in *Jag1^+/+^,* p-value =ns and 12.6±2.3 OPC cells per 150 µm in *Jag1^Ndr/Ndr^* compared to 12.2±2.3 in *Jag1^+/+^,* p-value =ns, mean ± SD. n=5 per genotype, with exception of P10 PC data (n=2); data are mean with standard deviation, scale bar represents 20 um; *p-value <0.05; **p-value <0.01; ***p-value <0.001, ****p-value <0.0001, unpaired t-test.

**Supp. Fig 2: The E14.5 *Jag1^Ndr/Ndr^* prosensory domain is smaller and exhibits reduced Notch activation. a,b )** Prosensory domain marker expression in the basal turn, including SOX2 (blue), JAG1 (red) and CDKN1B (green) for *Jag1^+/+^* mice (upper panels) and *Jag1^Ndr/Ndr^* mice (lower panels), indicating a smaller CDKN1B domain as well as a reduction in the SOX2-positive/JAG1-negative lateral prosensory domain (brackets), quantified in **b**). **c)** Notch target gene mRNA expression in the basal turn, showing reduced *HeyL, Hey1, Hes1* and *Hey2* expression in *Jag1^Ndr/Ndr^* mice, quantified in **d**) Number of dots for *Heyl* 80.0 ±48.4 in *Jag1^Ndr/Ndr^* compared to 173.3± 51.4 in *Jag1^+/+^* p-value <0.001, *Hey1* 334.5± 93.7 in *Jag1^Ndr/Ndr^* compared to 684.0± 103.1 in *Jag1^+/+^* p-value <0.01, *Hey2* 457.3 ±93.7 in *Jag1^Ndr/Ndr^* compared to 592.8± 55.5 in *Jag1^+/+^* p-value <0.01, *Hes1* in *Jag1^Ndr/Ndr^* 102.5± 33.4 compared to 254.3± 31.1 in *Jag1^+/+^,* p-value <0.001 n=5 per genotype; data is mean with standard deviation; scale bar represents 20 um; *p-value <0.05; **p-value <0.01; ***p-value <0.001, ****p-value <0.0001, unpaired t-test.

**Supp. Fig 3: mRNA expression of Notch components and genes with inducible Rbpj binding sites. a)** Violin plots showing the expression of Notch components in different cell types of the Organ of Corti, split per genotype. **b)** Dysregulation of genes with an inducible Rbpj binding sites^41^, in lateral *Jag1^Ndr/Ndr^* SCs. Red asterisks indicate significant upregulation (adj. p-value <0.05); black asterisks indicate potential upregulation (p-value <0.05).

**Supp. Fig 4: Subclustering and renormalization of HC populations. a)** UMAP projection of subsetted and renormalized HC population, split by genotype. **b**) Volcano plot showing downregulated genes (left, green) and upregulated genes (right, red) in *Jag1^Ndr/Ndr^* OHCs compared to *Jag^+/+^* OHCs, for the dataset shown in **a***.* **c**) Pseudotime analysis of OHC population shown in **a**, demonstrating that there is no significant difference in OHC pseudotime for *Jag1^Ndr/Ndr^* OHCs compared to *Jag^+/+^* OHCs. Data are mean with standard deviation; unpaired t-test.

**Supp. Fig 5: OHC-like cells are not present in *Jag1^+/Ndr^* mice. a)** HC phenotype of *Jag1^+/Ndr^* mice at E17.5 showing occasional supernumerary IHCs, but no OHC-like cells (MYO6, red). **b)** E17.5 quantification of IHCs, supernumerary IHCs (IHC +1), OHCs and OHC-like cells for all genotypes, 4.3±2.2 OHC-like cells per 100 µm in *Jag1^Ndr/Ndr^* compared to 0.0±0.0 in *Jag1^+/+^,* p-value <0.05, mean ±SD. **c)** *Jag1^+/Ndr^* phenotype at P50 showing OHC marker SLC26A5 (red) expression in the lateral OHC domain, but not outside of the lateral domain, indicating that there are no OHC-like cells in *Jag1^+/Ndr^* mice. n=5 per genotype; data are mean with standard deviation; scale bar represents 20um; *p-value <0.05; **p-value <0.01; ***p-value <0.001, ****p-value <0.0001, one-way ANOVA with Bonferroni correction.

**Supp. Fig 6: *Sez6l* and *Bmp2* expression in *Jag1^Ndr/Ndr^* OHC datasets and mRNA validation. a)** UMAP projection of unsupervised clustering of the renormalized HC subset, indicating an OHC population exclusively found in *Jag1^Ndr/Ndr^* OHCs (cluster3, blue). **b)** Featureplot showing expression of *Sez6l* and *Bmp2*, indicating high expression of Sez6l and Bmp2 in cluster 3. **c)** RNAscope for mRNA expression of *Sez6l* and *Bmp2* indicates expression of *Sez6l* and *Bmp2* in PCs in both *Jag1^Ndr/Ndr^* and *Jag^+/+^* mice, and absence of *Sez6l* and *Bmp2* in OHC-like cells. **d)** UMAP projection of OHCs subset with removal of a PC signature, showing similar clusters and lack of separation of additional clusters in *Jag1^Ndr/Ndr^* population, formerly present in **a**. n=3 for phenotypic mRNA analysis.

**Supp. Fig 7: Expression of immature and mature marker genes in OHC subset and pseudotime analysis**. **a)** UMAP showing expression of immature markers *Insm1* and *Blc11b* in OHC subset. **b)** UMAP showing expression of mature markers *Ikzf2* and *Slc26a5* in OHC subset. **c)** UMAP showing pseudotime. **d)** Pseudotime analysis of OHC population shown in **(a)**, demonstrating that there is no significant difference in OHC pseudotime for *Jag1^Ndr/Ndr^* OHCs compared to *Jag^+/+^* OHCs (4.0±2.9 in *Jag1^Ndr/Ndr^* compared to 4.6±3.1 in *Jag1^+/+^,* p-value =ns, mean ±SD)

**Supp. Fig 8: OHC-like cells do take up HC dye FM1-43. a)** FM1-43 uptake in *Jag^+/+^* HCs (green), indicating that IHCs and OHCs take up FM1-43 dye. **b)** FM1-43 uptake in *Jag1^Ndr/Ndr^* HCs (green), indicating that all HCs take up FM1-43 dye (left), including OHC-like cells (right, arrows). Scale bar represents 20um.

**Supplementary Table Legends**

**Table1**: Summary of Notch defective animal models reporting an inner ear phenotype, extended and updated table of previous published table1 in Brown et al, 2020 (doi: [10.3390/biom10030370](https://doi.org/10.3390%2Fbiom10030370))

**Table2:** Summary of hearing function Jag1 defective models

**Table3:** Marker genes for individual cell types for all Epcam^+^ cell populations (*Jag1^Ndr/Ndr^* and *Jag1^+/+^* combined).

**Table4:** Pseudo bulk analysis identified differently expressed genes for *Jag1^Ndr/Ndr^* versus *Jag1^+/+^,* pathway enrichment for differently expressed genes and previously reported Jag1 mediated genes.

**Table5:** Differentially expressed genes per cell type for *Jag1^Ndr/Ndr^* versus *Jag1^+/+^*

**Table6:** Pathway enrichment for differently expressed genes per cell type for *Jag1^Ndr/Ndr^* versus *Jag1^+/+^*

**Table7:** Marker genes for individual cell types for all Epcam^+^ cell populations for *Jag1^+/+^* dataset

**Table8:** Number of cells per cell type per genotype

**Table9:** OHC sub clustering analysis and PC signature removal

**Table10:** Primary antibodies and RNAscope probes
