## Supplement Figures for "Jag1 represses Notch activation in lateral supporting cells and inhibits an outer hair cell fate in the medial compartment of the developing cochlea"

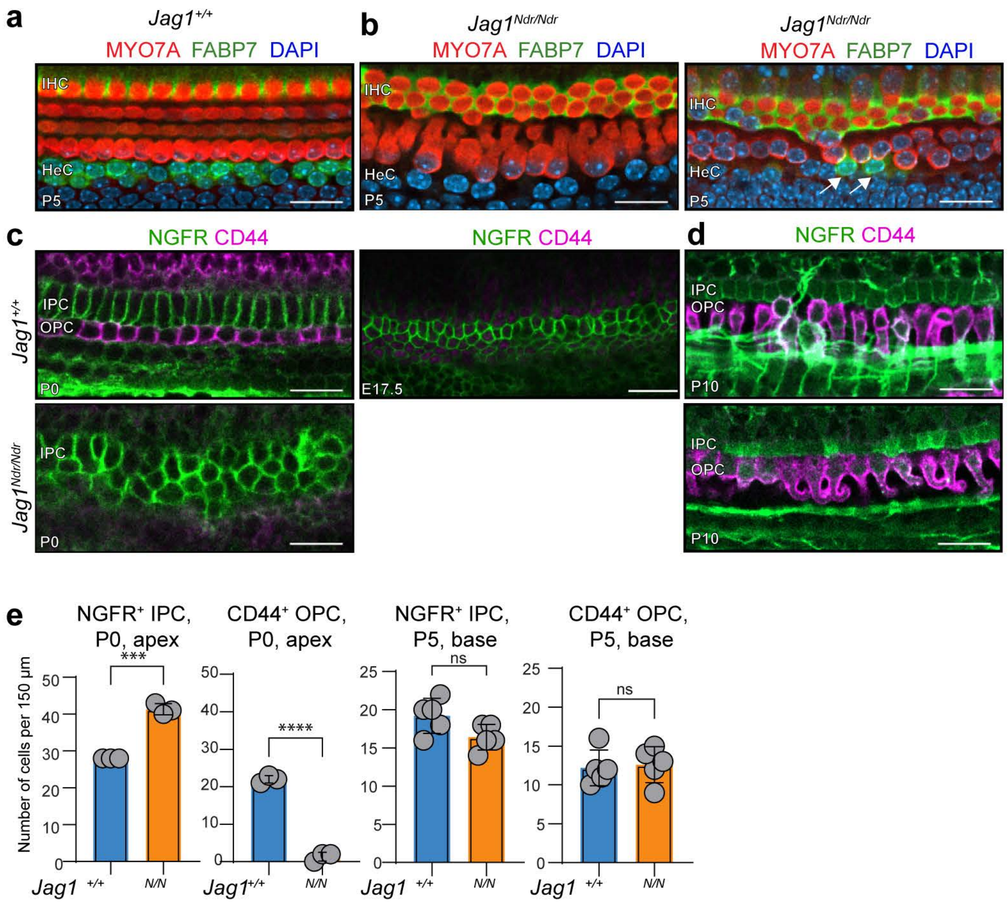


de Haan et al
Supplement Figure 1

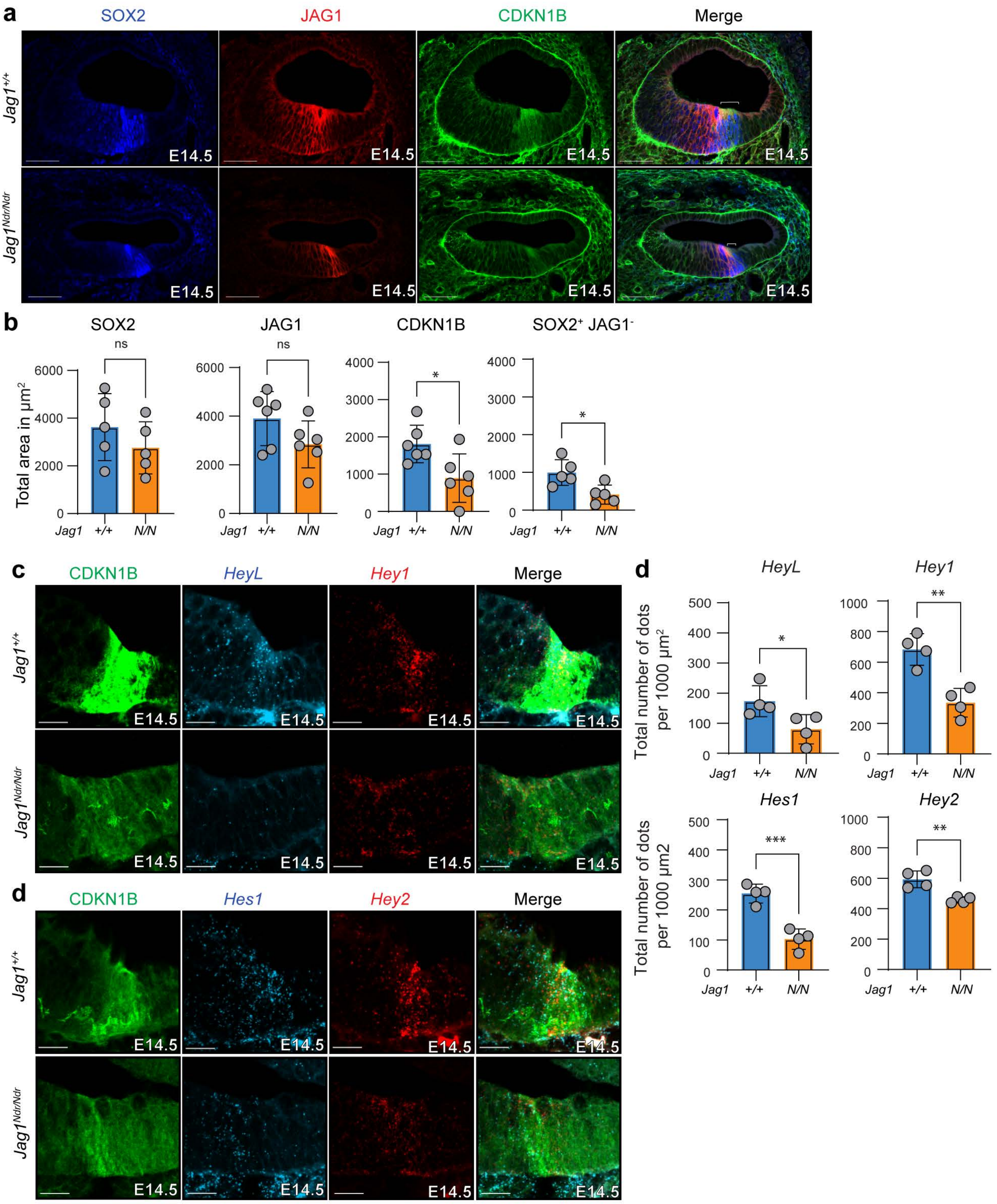


de Haan et al

Supplement Figure 2

**a** Notch component expression

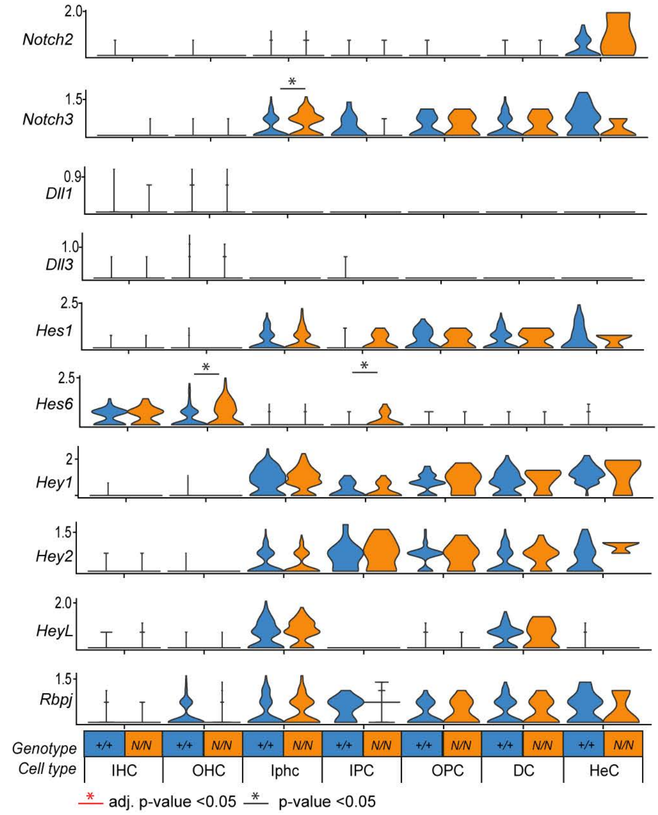


**b** Upregulated genes reported in Castel et al (doi: 10.1101/gad.211912.112)

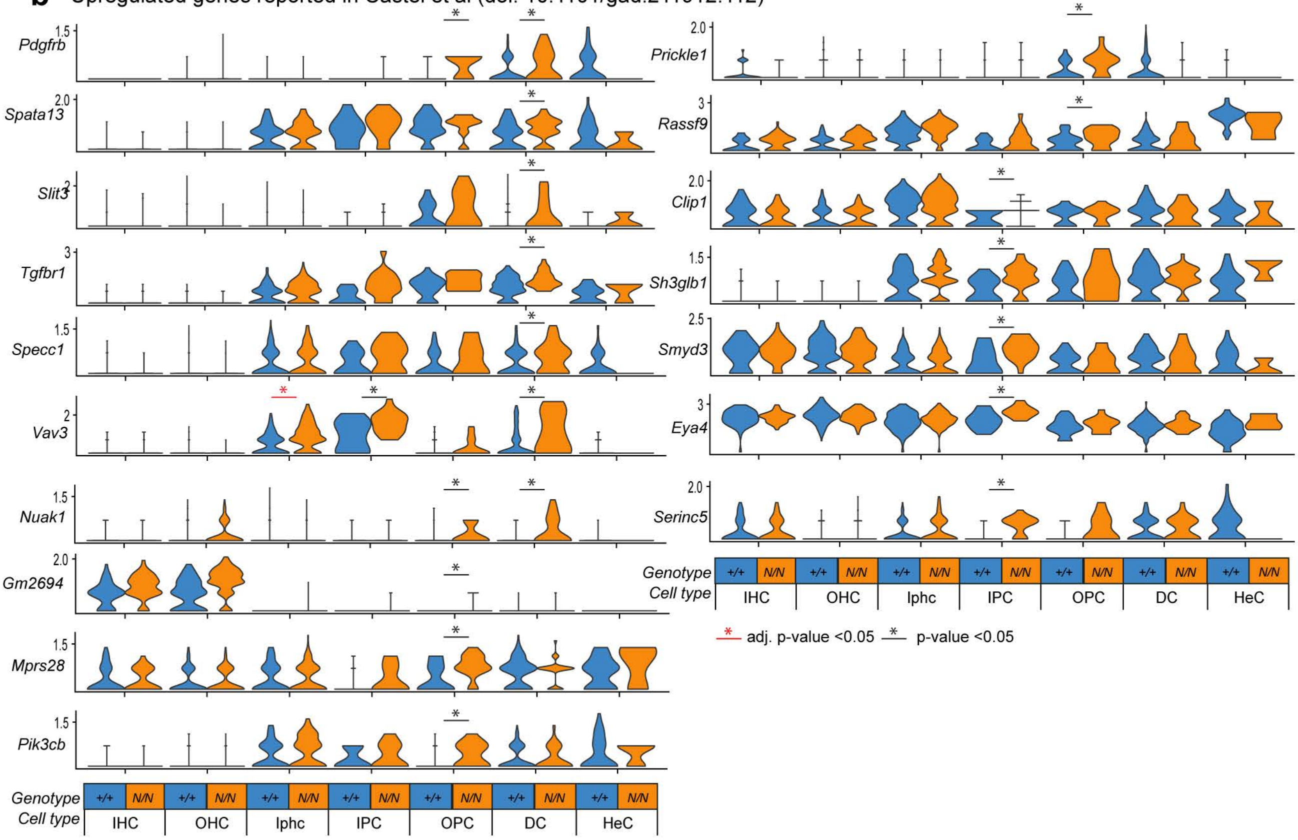


de Haan et al

Supplement Figure 3

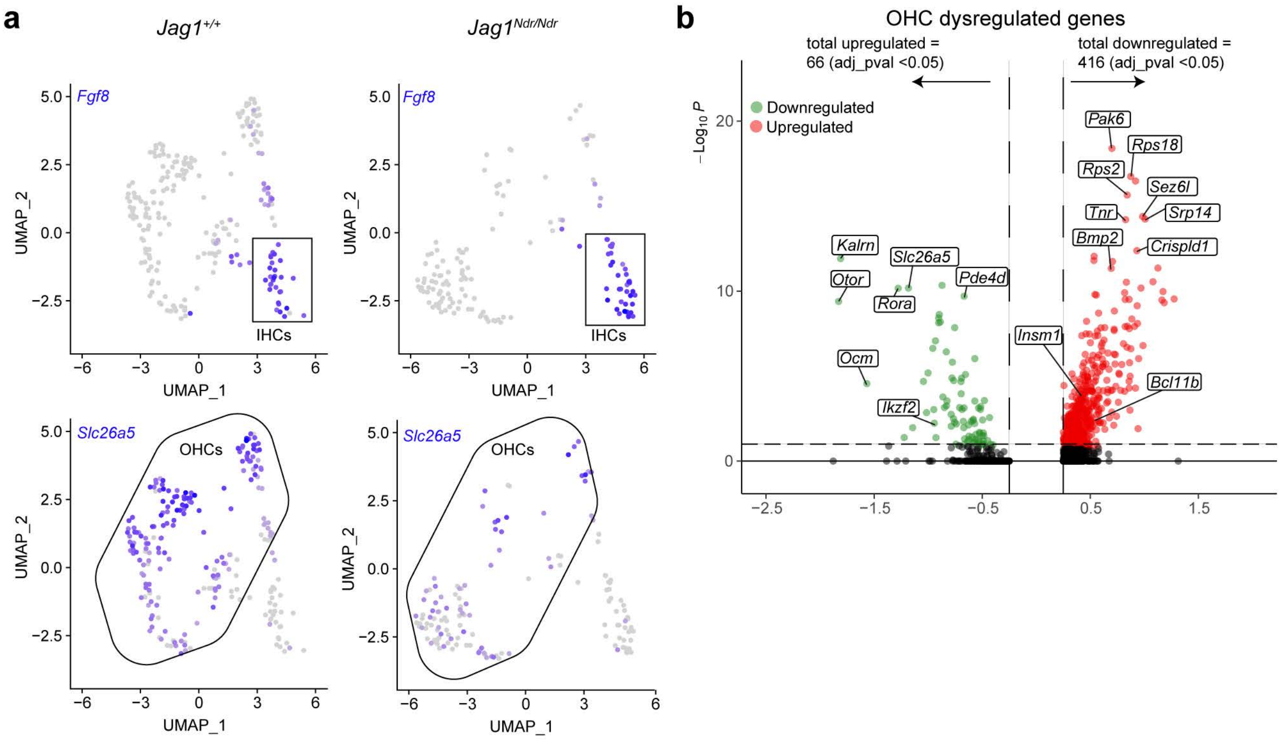


de Haan et al
Supplement Figure 4

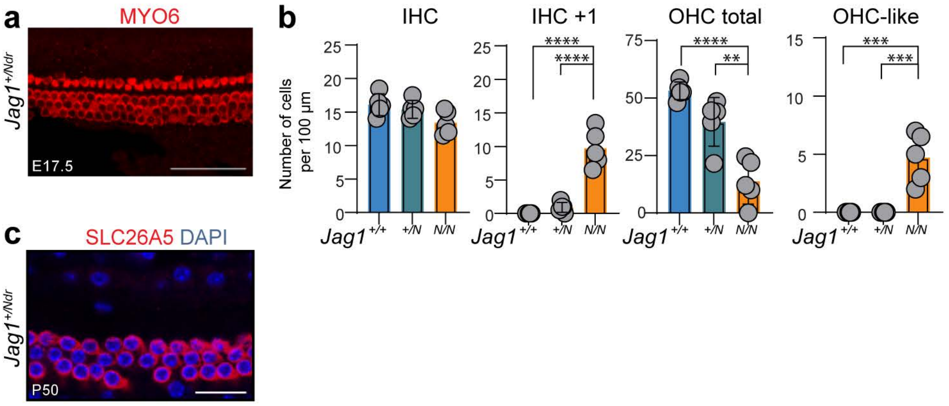


Supplement Figure 5
de Haan et al

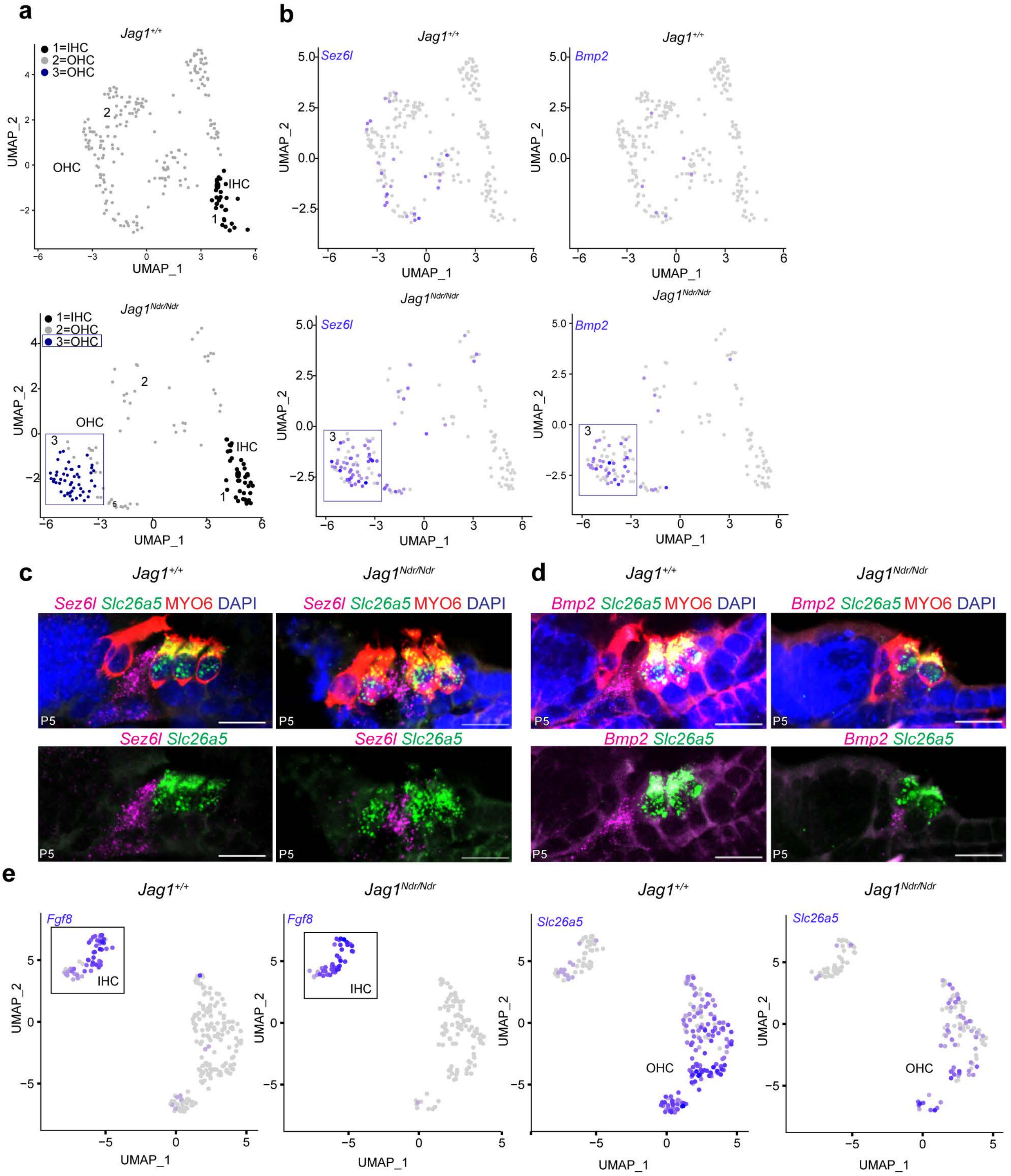


de Haan et al
Supplement Figure 6

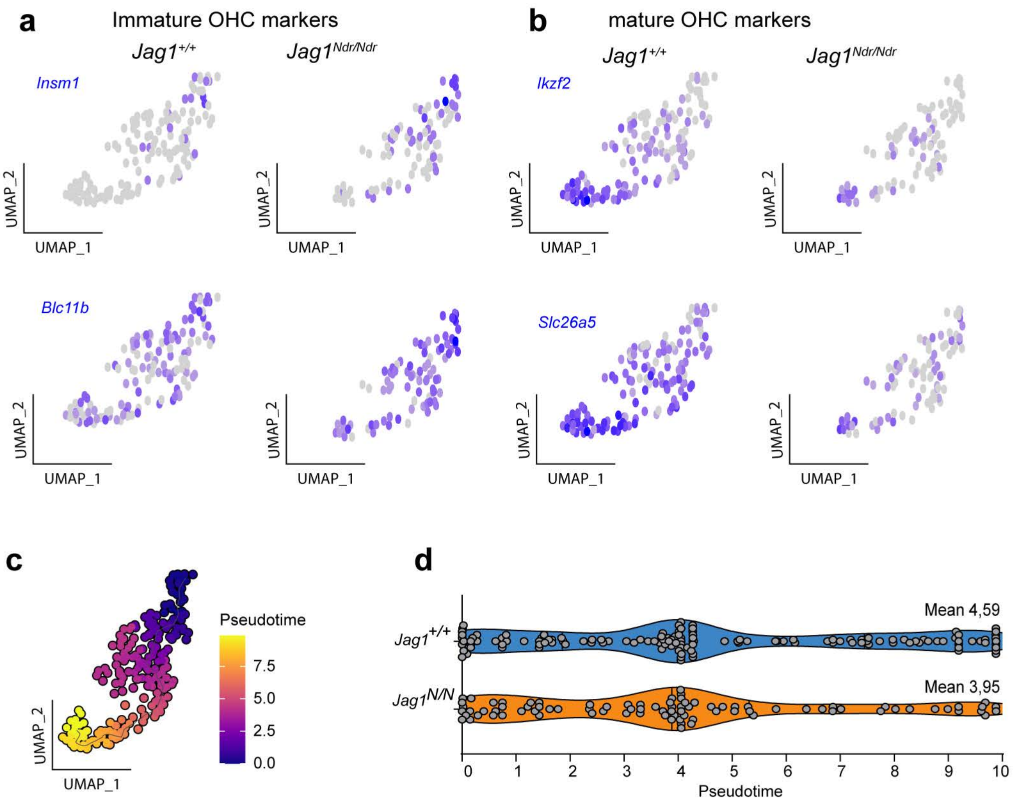


de Haan et al
Supplement Figure 7

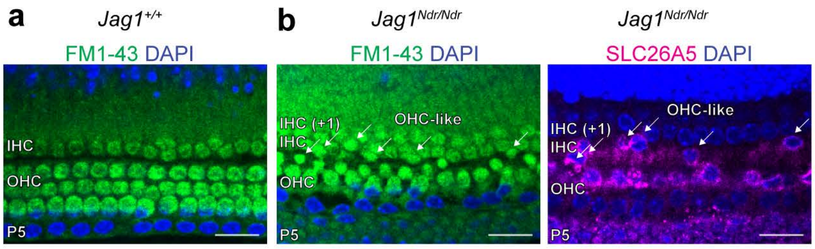


de Haan et al

Supplement Figure 8
