## Supplementary figures and images for "Jag1 represses Notch activation in lateral supporting cells and inhibits an outer hair cell fate in the medial compartment of the developing cochlea"

### Graphical abstract

#### Graphical abstract

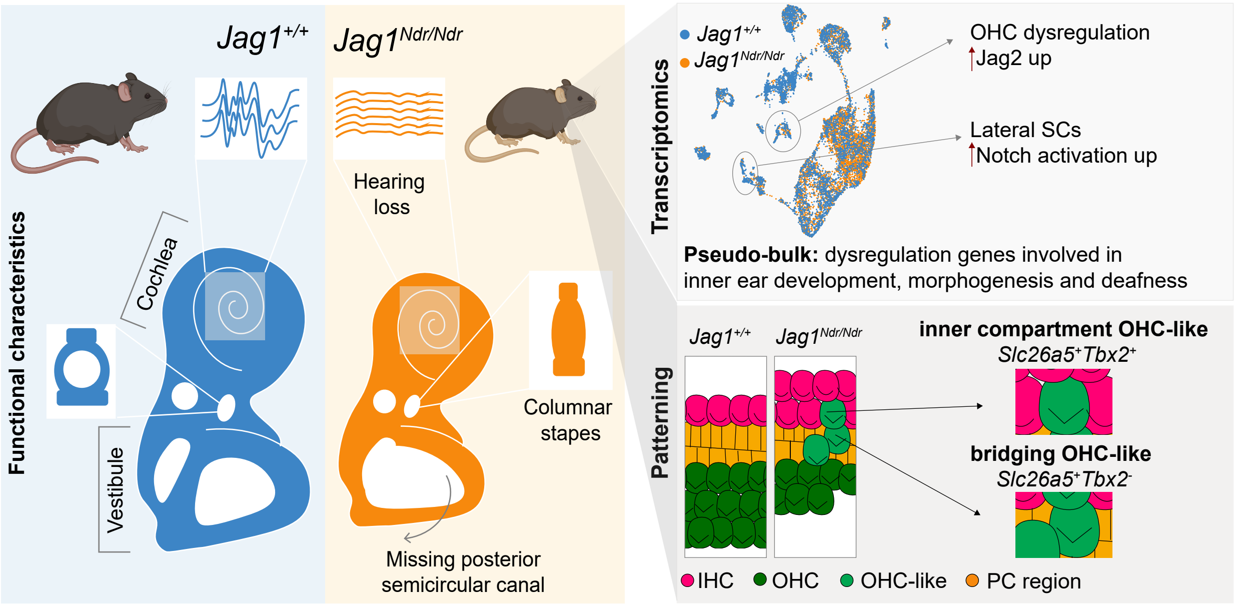
